# The ESKAPE mobilome contributes to the spread of antimicrobial resistance and CRISPR-mediated conflict between mobile genetic elements

**DOI:** 10.1101/2022.01.03.474784

**Authors:** João Botelho, Adrian Cazares, Hinrich Schulenburg

**Affiliations:** Antibiotic Resistance Evolution Group, Max Planck Institute for Evolutionary Biology, Plön, Germany; Department of Evolutionary Ecology and Genetics, Zoological Institute, Christian Albrechts University, Kiel, Germany; EMBL’s European Bioinformatics Institute (EMBL-EBI), Wellcome Genome Campus, Cambridge, United Kingdom; Wellcome Sanger Institute, Wellcome Genome Campus, Cambridge, United Kingdom

## Abstract

Mobile genetic elements (MGEs) mediate the shuffling of genes among organisms. They contribute to the spread of virulence and antibiotic resistance genes in human pathogens, such as the particularly problematic group of ESKAPE pathogens, including *Enterococcus faecium*, *Staphylococcus aureus*, *Klebsiella pneumoniae*, *Acinetobacter baumannii*, *Pseudomonas aeruginosa*, and *Enterobacter* sp. Here, we performed the first systematic analysis of MGEs, including plasmids, prophages, and integrative and conjugative/mobilizable elements (ICEs/IMEs), in the ESKAPE pathogens. We characterized over 1700 complete ESKAPE genomes and found that different MGE types are asymmetrically distributed across these pathogens. While some MGEs are capable of exchanging DNA beyond the genus (and phylum) barrier, most horizontal gene transfer (HGT) events are restricted by phylum or genus. We show that the MGEs proteome is involved in diverse functional processes and distinguish widespread proteins within the ESKAPE context. Moreover, anti-CRISPRs and antimicrobial resistance (AMR) genes are overrepresented in the ESKAPE mobilome. Our results also underscore species-specific trends shaping the number of MGEs, AMR, and virulence genes across pairs of conspecific ESKAPE genomes with and without CRISPR-Cas systems. Finally, we found that CRISPR targets vary according to MGE type: while plasmid and prophage CRISPRs almost exclusively target other plasmids and prophages, respectively, ICEs/IME CRISPRs preferentially target prophages. Overall, our study highlights the general importance of the ESKAPE mobilome in contributing to the spread of AMR and mediating conflict among MGEs.

## Introduction

Mobile genetic elements (MGEs) are DNA entities that are capable of capturing and shuffling genes intra- and intercellularly [1]. Coevolution of bacterial hosts with these MGEs has driven the evolution of complexity [2]. Movement within the genome is often mediated by specific MGEs, such as insertion sequences and transposons [3]. Others like plasmids, prophages, and integrative and conjugative/mobilizable elements (ICEs/IMEs) are key vectors for intercellular mobility, being responsible for a large fraction of the variability observed between bacterial species [4–7]. Bacteria undergo extensive horizontal gene transfer (HGT), and some estimates suggest that more than 80% of bacterial genes were horizontally transferred at some point in their evolutionary history [8]. These events are largely shaped by ecological niches, by the difference in the GC content between pairs of bacteria exchanging material, and by phylogenetic barriers [9–11]. Network-based methods are useful to trace HGT events and recover shared content between bacterial genomes [11,12], and have been recently applied to explore the population structure of thousands of plasmids [13,14]. Even though this approach has been useful to explore population structure of plasmids, the study of potential HGT events involving other MGEs (such as prophages and ICEs/IMEs) is largely unexplored. Moreover, only a few studies have used network-based approaches to explore the co-evolutionary dynamics of different MGE types [15–17].

MGEs carry non-essential genes that can provide their bacterial host with adaptive traits and alter their fitness, such as antimicrobial resistance (AMR) and virulence genes [18,19]. These elements apply a myriad of ecological and evolutionary strategies to promote their own replication and transmission, which allow them to persist even in the absence of positive selection for the beneficial genes they carry [20,21]. As a predicted immune response against invading MGEs, bacteria have evolved different, often complex, mechanisms such as restriction-modification and clustered, regularly interspaced short palindromic repeats (CRISPR) and CRISPR-associated (Cas) genes [22]. These systems are usually clustered in ‘defence islands’ and are widespread in bacteria and archaea [23,24]. Spacer sequences, derived from fragments of invading MGEs, are continuously incorporated at CRISPR loci, enabling partial reconstruction of recent HGT events. Curtailment of HGT through CRISPR-Cas immunity can be favourable when dealing with obligate parasites, but can be detrimental by hindering the acquisition of genetic novelty and/or beneficial traits carried by MGEs. Invasion by these MGEs can, however, be associated with fitness costs that may lead to selection against carriage [5,25]. Hence, bacteria often face a trade-off between immunity and acquisition of novel elements, which favour adaptation to different ecological niches and stressors, such as antibiotic pressure. MGEs can be equipped with inhibitors of CRISPR-Cas systems, called anti-CRISPR (Acr) proteins, which have been reported mostly in prophages [26–28]. Recently, Acr proteins were identified in non-phage MGEs, including plasmids and ICEs [29].

Bacterial pathogens belonging to the ESKAPE panel consist of five species (*Enterococcus faecium*, *Staphylococcus aureus*, *Klebsiella pneumoniae*, *Acinetobacter baumannii*, and *Pseudomonas aeruginosa*) and one genus (*Enterobacter* sp.) [30,31]. These pathogens are frequently involved in problematic nosocomial infections, due to their multi-drug resistance and/or invasive phenotypes [32–37]. The WHO recently published a list of pathogens for which new antibiotic development is urgently required, and the ESKAPE pathogens were designated “priority status” [38]. AMR and virulence genes are broadly distributed in plasmids across the ESKAPE pathogens [19,32], and also in ICEs [39,40]. Recently, CRISPR-Cas systems have been identified in plasmids and ICEs from several bacterial species (including representatives of the ESKAPE pathogens), and may be involved in conflict between MGEs [41–43].

In this study, we performed the first systematic analysis focusing on the ESKAPE pathogens mobilome. We asked i) how prevalent are different MGEs (prophages, ICEs/IMEs, and plasmids) across the ESKAPE pathogens; ii) how broad or constrained is the combined MGEs’ network; iii) which traits are overrepresented in these MGEs, and if AMR and virulence genes are differently distributed in pairs of conspecific ESKAPE pathogens with and without CRISPR-Cas systems; iv) whether the CRISPR spacers have a targeting bias towards different MGE types. We found that plasmids, ICEs/IMEs, and prophages are unequally distributed across these pathogens, and found signatures of DNA sharing events between different species. Uncovering the structure of MGEs and masked (i.e., MGE-free) genomes allowed us to discover an overrepresentation of AMR genes and anti-CRISPRs in the ESKAPE mobilome. Our results also unveiled ESKAPE-specific trends of MGEs, AMR, and virulence genes promoted by the presence of CRISPR-Cas systems. Finally, our work shows that CRISPR spacers found on prophages, ICEs/IMEs, and plasmids across the ESKAPE pathogens have different targeting biases.

## Results

### 1. MGEs are unevenly distributed among the ESKAPE pathogens

We downloaded 1782 ESKAPE complete genomes from NCBI’s RefSeq database. To correct for species taxonomy, genomes with <95% average nucleotide identity (ANI) were removed for each ESKAPE species (**Table S1)**. Since this parameter is only applied for species delineation, we also built a phylogenomic tree with *Enterobacter* sp. genomes and type strains belonging to the *Enterobacteriaceae* family (**Figure S1**). Our curated dataset included 1746 complete genomes, belonging to multiple MLST profiles (**Table S2**). We found a total of 21478 MGEs, including 16153 prophages, 2685 ICEs/IMEs, and 2640 plasmids (**Figures 1A and 1D**). We confirmed that plasmids were abundant in *K. pneumoniae* (∼3 plasmids/genome). Actually, these replicons were prevalent in every ESKAPE except *P. aeruginosa* and *S. aureus* (**Figure 1A**). The majority of plasmids carried a relaxase (62.5%, 1651/2640), and were classified as mobilizable (either self-conjugative or not) [44].

**Figure 1.**
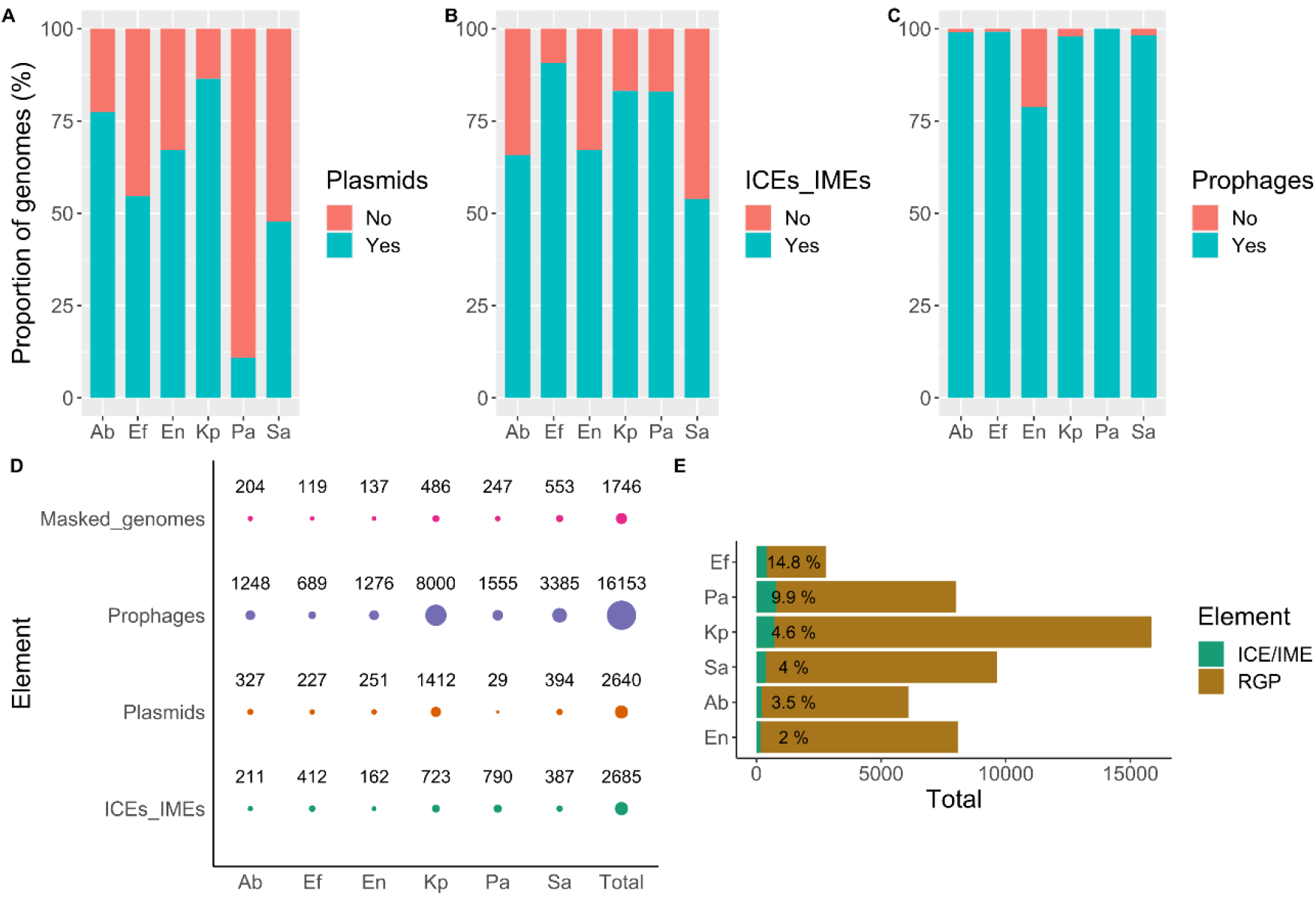
Distribution of MGEs across the ESKAPE pathogens. Proportion of genomes carrying at least one **A)** plasmid, **B)** ICE/IME, or **C)** prophage. **D)** Total number of MGEs and of considered masked genomes per ESKAPE pathogen. Size of the circles is proportional to the number of identified elements. **E)** Total number of RGPs and ICEs/IMEs per ESKAPE. The size of the green bars is proportional to the total number of ICEs/IMEs identified per ESKAPE pathogen, and the relative number of ICEs/IMEs per RGPs is shown in percentage next to the green bars. Bars are sorted according to the relative number of ICEs/IMEs per RGPs. Ab, *A. baumannii*; Ef, *E. faecium*; En, *Enterobacter* sp.; Kp, *K. pneumoniae*; Pa, *P. aeruginosa*; Sa, *S. aureus*.

To look for regions of genome plasticity (RGPs) exclusively integrated in the chromosome, we used the 1746 chromosomal replicons to generate plasmid-free pangenomes for each ESKAPE taxon. We identified a total of 50482 plasmid-free RGPs in chromosomal replicons (**Figure 1E**). Of these, 2685 were classified as ICEs/IMEs due to the presence of relaxase and integrase domains (**Figures 1D and 1E**). At least one ICE/IME was detected in >50% of genomes for all ESKAPE pathogens, and was abundant in *E. faecium* and *P. aeruginosa* (∼3 elements/genome) (**Figures 1B and 1D**). After masking the ICEs/IMEs identified in the ESKAPE chromosomes, we performed a search for prophages. These elements were the most abundant MGE type found in the ESKAPE collection. Additionally, prophages were significantly more prevalent than ICEs/IMEs and plasmids across all ESKAPE pathogens (**Figure S2**).

When looking into the presence/absence combination of co-occurring MGEs across the ESKAPE pathogens, we noticed that the most frequent combination involved the presence of the three MGEs (in 717 out of the 1746 genomes, **Table S2**). We noticed that the majority of the strains with the three MGEs co-occurring in the same genome belonged to *K. pneumoniae* (340/717). Our results show that different MGEs are asymmetrically distributed across the ESKAPE pathogens, with *K. pneumoniae* genomes taking the lead for the co-occurrence of ICEs/IMEs, plasmids, and prophages (**Figure 1**).

### 2. DNA sharing between different vectors is common across the ESKAPE mobilome

MGEs tend to have a GC content lower than the one from its host [45–47]. However, how conserved is this difference when considering different MGE types is poorly studied. We confirmed that for most MGE/ESKAPE pairs, the arithmetic mean GC content of the different MGEs is significantly lower when compared to masked genomes across the ESKAPE pathogens (**Figure S3A**, p-value < 2.2e-16). With the exception of *S. aureus*, we observed that plasmids across the ESKAPE pathogens have a wider variation in their size, when compared with ICEs/IMEs and prophages (**Figure S3B**). Across all ESKAPE pathogens, we observed a weak positive correlation between the ICEs/IMEs and plasmids’ GC content and sequence length (R = 0.38 and 0.35, respectively, p<2.2e-16), and a weak negative correlation between the prophages’ GC content and sequence length (R = −0.15, p<2.2e-16, **Figure S4**).

Given the presence of highly similar MGEs in our dataset, we dereplicated the 21478 elements found here into a representative set of 10339 MGEs. We then used an alignment-free sequence similarity comparison of the ESKAPE mobilome to infer an undirected network (**Figures 2A and 2B**). The sparse network assigned 97.8% (10110/10339) of the MGEs into 87 clusters. The majority of MGE pairs shared little similarity, with a Jaccard Index (JI) value below 0.25 (**Figure S5A**), in accordance with the high diversity frequently observed across MGEs. The network revealed clear structural differentiation, where the majority of the smaller clusters were homogeneous for a given ESKAPE/MGE pair (**Figures 2A and 2B**). The absence of pairwise distance similarities with intermediate JI (**Figure S5A**) helps to explain this clustering in discrete groups, instead of a continuous genetic structure. However, the two largest clusters challenge interspecies and MGE type barriers and correspond to multiple MGEs with the four Proteobacteria representatives in the first (i.e., *K. pneumoniae*, *Enterobacter* sp., *P. aeruginosa*, and *A. baumannii*), and *S. aureus* and *E. faecium* in the second cluster. MGEs within these promiscuous clusters tend to be more dissimilar than those assigned to densely connected and ESKAPE/MGE pair-restricted clusters. To plot this network, we used as a threshold the mean value (0.0537361) of the estimated pairwise distances between the 10110 MGEs identified in this study. In parallel, we built a network using the masked genomes as nodes connected by edges indicating pairwise distances, and we exclusively observed species-specific clustering (**Figures S5B and S6**), which is in agreement with cohesive evolutionary forces that maintain the genomic coherence of bacterial species [48,49]. We observed that DNA sharing events between different MGE/ESKAPE pairs are pervasive (**Figure S7**). We also observed cross-phylum interactions, between different MGEs across the ESKAPE pathogens. Altogether, our results highlight that while some elements are capable of exchanging DNA beyond the genus (and phylum) barrier, for the majority of clusters host similarity and MGE type restrain DNA sharing between different ESKAPE MGEs.

**Figure 2.**
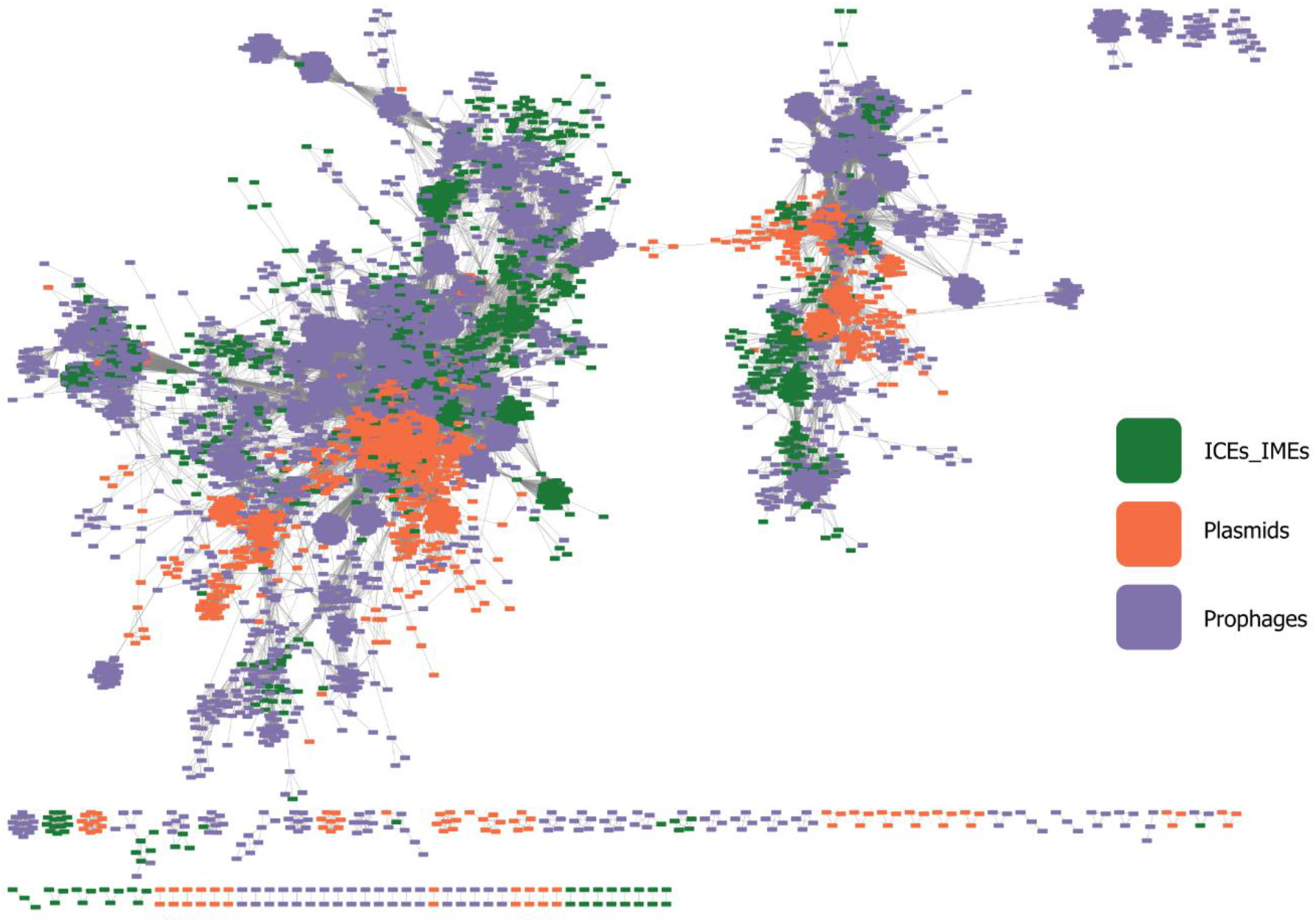
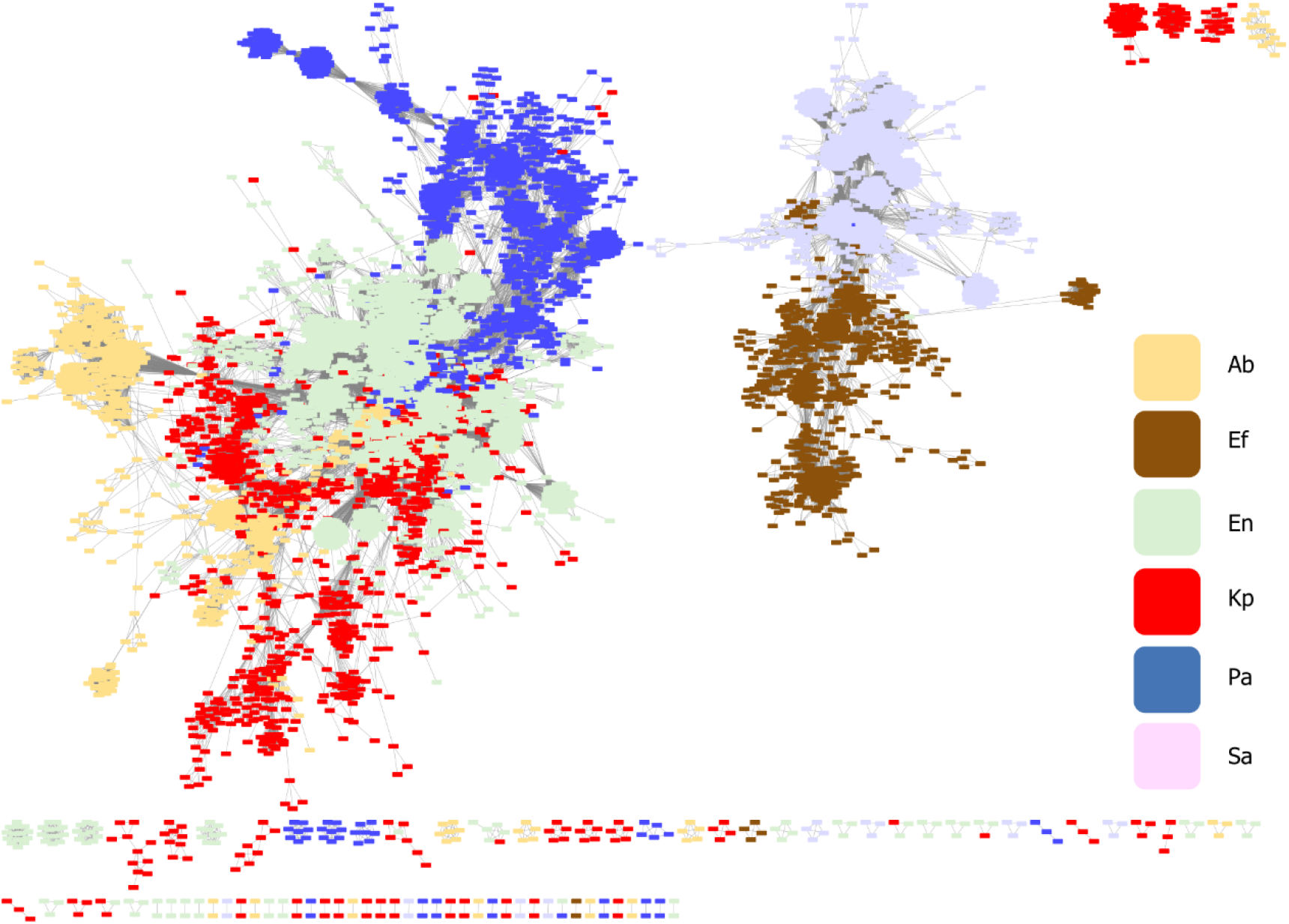
Network of clustered MGEs, using the mean Jaccard index as a threshold. Network grouped by **A)** MGE type; **B)** ESKAPE pathogen. Each MGE is represented by a node, connected by edges according to the pairwise distances between all MGE pairs. The network has a clustering coefficient of 0.781, a density of 0.014, a centralization of 0.065, and a heterogeneity of 0.959. Ab, *A. baumannii*; Ef, *E. faecium*; En, *Enterobacter* sp.; Kp, *K. pneumoniae*; Pa, *P. aeruginosa*; Sa, *S. aureus*.

### 3. The proteome of the ESKAPE mobilome is highly diverse in sequence and functions

To gain functional insights into the proteome of the ESKAPE mobilome, we investigated the diversity of clusters of orthologous groups (COGs) encoded by MGEs identified in this study. COGs are protein sets conserved across lineages that typically share function and are therefore used for systematic function prediction in poorly characterised genomes [50]. COGs have been assigned to curated and uniform functional categories, thus enabling the comparison of their distribution amongst genomes [51]. We distinguished 2761 different COGs in the ESKAPE MGEs (**Figure S8**). These clusters encompass most functional categories reported in the COGs scheme (**Figure 3**), thereby highlighting the diversity of functions associated with the ESKAPE mobilome. ICEs, plasmids, and prophages contain 148, 164, and 794 unique COGs, respectively, consistent with their distinctive features as mobile elements (**Figure S8**). However, they also share 921 COGs (∼33%), indicative of a common pool of proteins and functions carried by MGEs in ESKAPE pathogens. COGs present in only two of the three MGE types were also identified, with prophages and plasmids sharing more COGs than other pairs (**Figure S8**).

**Figure 3.**
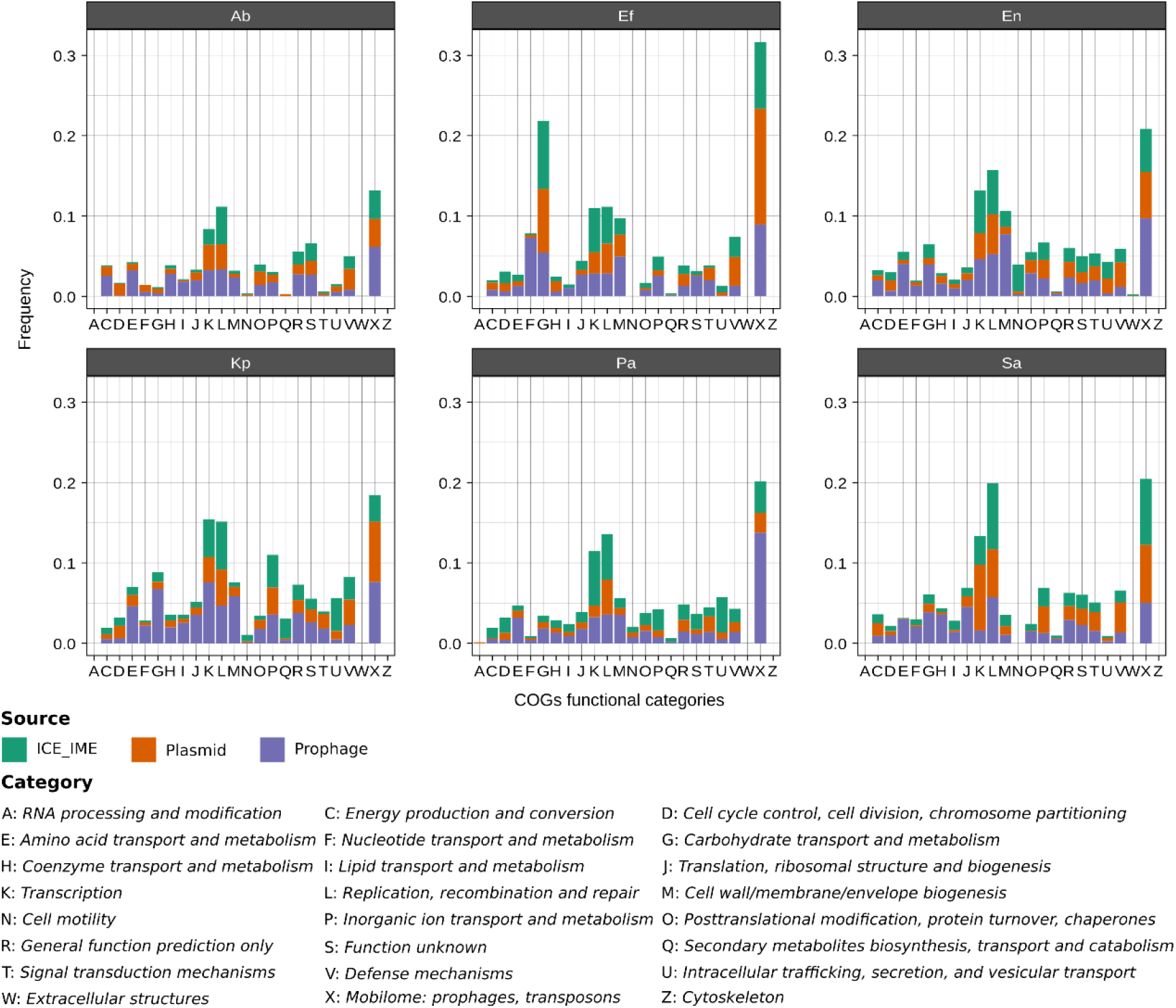
Relative contribution of ESKAPE MGEs proteome to COG functional categories. The barplots in the figure are split into facets corresponding to the different ESKAPE pathogens. The relative frequency of proteins in the different COG functional categories was calculated separately for each MGE type by dividing the number of proteins belonging to a given category by the total number of proteins observed in the corresponding MGE/ESKAPE pair. Hence, the bars illustrate the incidence of proteins of a given functional category in the proteomes of the different MGE types per ESKAPE. The COG functional categories are indicated on the X axis and described at the bottom of the figure.

From 36 to 55% of proteins in the different MGE proteomes were assigned to a COG (see methods). Interestingly, we detected some variation in the relative contribution of MGE proteomes to different COG functional categories among the ESKAPE pathogens (**Figure 3**). For example, proteins associated with “Carbohydrate transport and metabolism” (category G) were more frequent in MGE proteomes of *E. faecium* than in other ESKAPE (**Figure 3**). The relative frequency of proteins in the “Cell wall/membrane/envelope biogenesis” category (M) also varied noticeably across MGE/ESKAPE pairs. On the other hand, proteins of the “Transcription” and “Replication, recombination and repair” categories (K and L, respectively) were among the most frequent in the MGE proteomes of all ESKAPE. As expected, proteins assigned to the COGs mobilome category (X) dominated all the MGE proteomes.

To explore the diversity of the ESKAPE MGEs proteome further, we clustered their 943246 proteins based on sequence similarity, resulting in 72247 groups (**Table S3**). Around 69% of the representatives of these protein groups were assigned the tag “hypothetical protein” by prokka, underlining the large proportion of uncharacterised proteins encoded by ESKAPE MGEs. Among the representatives with an assigned function, transposases and integrases were the most frequent protein product (2290 cluster representatives) (**Table S3**). Recombinase, repressor and resistance, were also common terms across the representative products with more than 200 occurrences each; the latter being mostly associated with metal or drug resistance.

We then looked for protein clusters widespread in MGE proteomes, i.e. those observed in the three MGE types or in at least three of the ESKAPE pathogens. Our search resulted in the identification of 1421 protein clusters widespread across MGEs and 426 present in at least three different ESKAPE (**Table S3**), with 187 clusters identified in common between the two widespread categories. Although hypothetical proteins dominated both widespread categories (55% of MGEs and 50% of ESKAPE widespread protein clusters), various protein clusters with functions associated with transposition and antimicrobial resistance were also identified (**Table S3**). Hierarchical clustering of the MGE/ESKAPE pairs and widespread protein clusters based on the distribution and relative frequency of the latter uncovered structured patterns of sharing (**Figure 4**). For example, we detected a component of ESKAPE-widespread protein clusters present in plasmids of *Enterobacter* sp., *K. pneumoniae*, and ICEs/IMEs of *P. aeruginosa*. When it comes to protein clusters present in different MGE types, we observed clusters predominantly occurring in ICEs/IMEs and phages of *A. baumannii* and *E. faecium*. Overall, the distribution of ESKAPE widespread proteins clustered MGE/ESKAPE pairs by MGE type (**Figure 4**). The clustering observed from the distribution of MGE widespread proteins was more intricate, with only a couple of clusters featuring the same ESKAPE pathogen. Altogether, our results show that more than seventy thousand protein clusters, representing nearly one million sequences, are linked to the mobilome component of the ESKAPE pangenomes. These proteins are involved in a broad range of functional categories; frequently in transcription, replication and recombination. Only ∼2.3% of protein clusters are widespread within the ESKAPE MGEs context but they feature complex distribution and frequency patterns.

**Figure 4.**
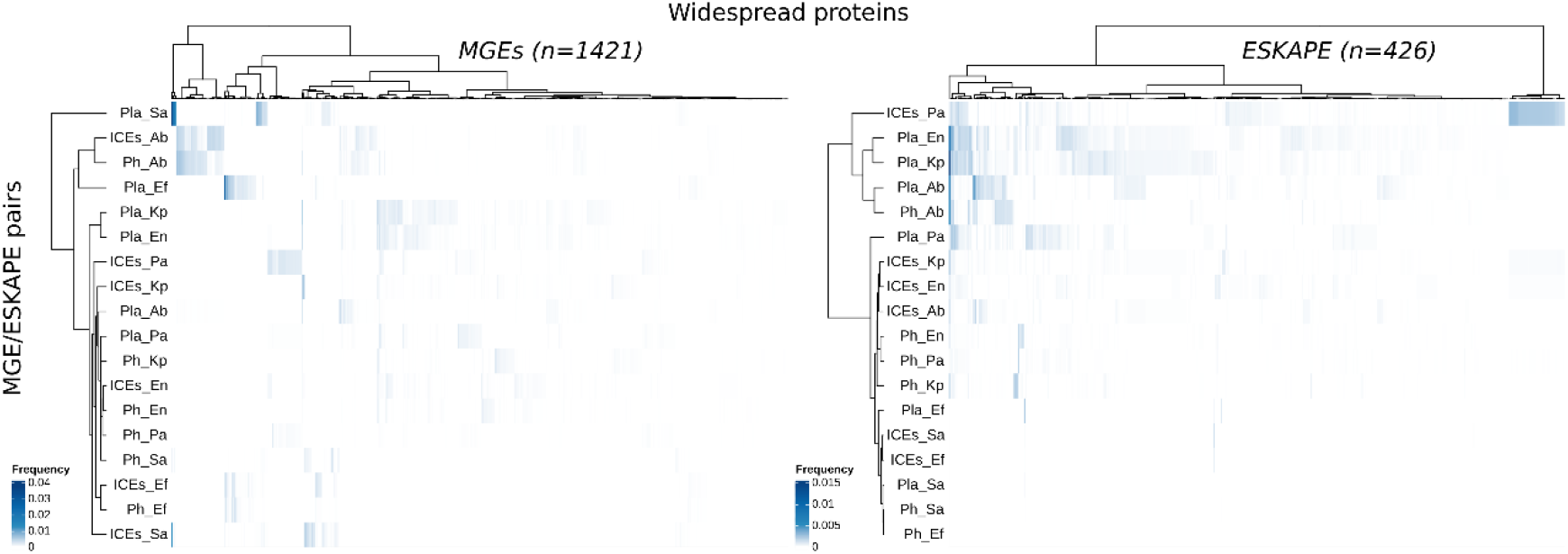
Distribution of widespread protein clusters across MGE/ESKAPE pairs. The heatmaps show the distribution and relative frequency of protein clusters (columns) identified as widespread in either MGEs (i.e. present at least once in the three MGE types; left side) or ESKAPE pathogens (i.e. present in at least three different ESKAPE; right side). The number of protein clusters represented in the heatmaps is shown in parenthesis. MGE/ESKAPE pairs are indicated on the left side of the heatmaps. The relative frequency of the protein clusters in the different MGE/ESKAPE pairs was calculated by dividing the number of occurrences identified by the total number of proteins observed in a given pair. The trees displayed at the top and left side of the heatmaps illustrate the hierarchical clustering of protein clusters and MGE/ESKAPE pairs using the ward.D method with euclidean distance. A list of the protein clusters, their products and relative frequency values is provided in **Table S3**.

### 4. AMR genes are overrepresented in the ESKAPE mobilome

In order to explore the AMR repertoire of the ESKAPE mobilome, we only focused on genes that are horizontally transferred, such as beta-lactamases and aminoglycoside-modifying enzymes (that lead to antimicrobial inactivation), and those that promote target site modification (such as rRNA methyltransferases, the *vanA* and *vanB* gene clusters, and the staphylococcal cassette chromosome *mec*). We observed that AMR genes are broadly distributed in plasmids across the ESKAPE pathogens. Even though the total number of prophages far outnumber that of plasmids in our collection (**Figure 1**), the absolute count of AMR genes in plasmids is greater than that observed in prophages (6068 versus 1845, respectively) (**Figure S9**). Interestingly, most AMR genes in plasmids and prophages are found in *K. pneumoniae*, whereas *P. aeruginosa* carries the majority of these genes within ICEs/IMEs (**Table S4**). All ESKAPE pathogens have a large proportion of AMR-carrying plasmids (>35% of plasmids across the different ESKAPE carry at least one AMR gene), while a high proportion of AMR-harbouring ICEs (>25%) was only observed for *S. aureus* and *P. aeruginosa* (**Figures S10A and S11B**). As previously reported [52], we observed that AMR genes are rarely found in prophages (**Figure S10C**). As expected from the vast repertoire of MGEs present in *K. pneumoniae* (**Figure 1D**), this species presented a wider selection of different AMR genes. Some AMR genes were restricted to specific ESKAPE pathogens, while others were more promiscuous. For example, different beta-lactamases (*bla* genes) were prevalent among proteobacterial representatives of the ESKAPE pathogens, but were mostly absent from *S. aureus* and *E. faecium* (**Figure S9** and **Table S4**). The only exception was the *blaZ* gene, which was exclusively identified in plasmids, prophages, and ICEs/IMEs from *S. aureus*. This gene is typically embedded within the SCCmec elements of this species and may have been acquired from distantly related non-*Staphylococcus* species [53]. Genes encoding resistance to aminoglycosides (*aac*, *ant*, and *aph* genes), chloramphenicol (*cat* genes), trimethoprim (*dfr* genes), and tetracyclines (*tet* genes) were found in all representatives of the ESKAPE pathogens. Genes involved in resistance to vancomycin (the *vanHAX* and *vanHBX* gene clusters) were exclusively found in *S. aureus* and *E. faecium*, while genes coding resistance for quinolones (*qnr* genes) and colistin (*mcr* genes) were only found in the proteobacterial representatives (**Figure S9** and **Table S4**).

We next assessed the distribution of virulence genes. These genes are broadly distributed in prophages and ICEs/IMEs across the ESKAPE pathogens **(Figure S11** and **Table S5**). In fact, we identified no virulence genes in *E. faecium* plasmids, and only 0.6% of *A. baumannii* plasmids carry these genes. More than 25% ICEs/IMEs in *S. aureus*, *K. pneumoniae*, and *E. faecium* carried at least one virulence gene (**Figures S10C and S10D**). *P. aeruginosa* is the ESKAPE pathogen carrying a wider variety of virulence genes in these MGEs, mostly on prophages. Polyketide synthesis loci *ybt* and *clb* encoding the iron-scavenging siderophore yersiniabactin and genotoxin colibactin, respectively, were solely identified in *Enterobacteriaceae* representatives of the ESKAPE pathogens (i.e., *K. pneumoniae* and *Enterobacter* sp.). These virulence loci were mostly present in ICEs/IMEs, as previously reported [54], but we also found these genes on plasmids and prophages (**Figure S11** and **Table S5**). Interestingly, *S. aureus* was the ESKAPE pathogen with a higher proportion of both plasmids and ICEs/IMEs carrying at least one AMR or virulence genes (**Figure S10**).

Since chromosomes are substantially larger than MGEs and consequentially have more genes, we corrected the prevalence of AMR and virulence genes to the total number of genes present in MGEs and masked genomes across the different ESKAPE pathogens. Overall, we noticed that AMR genes were largely overrepresented in MGEs (∼5x), when compared with masked genomes (**Figure 5**). On the other hand, virulence genes were ∼2x more likely to be located on masked genomes. Taken together, our results show that plasmids are important vectors for AMR genes across the ESKAPE pathogens, while ICEs/IMEs and prophages play a more important role for the distribution of virulence genes. When compared with masked genomes, we confirmed that AMR genes are preferentially distributed in the ESKAPE mobilome.

**Figure 5.**
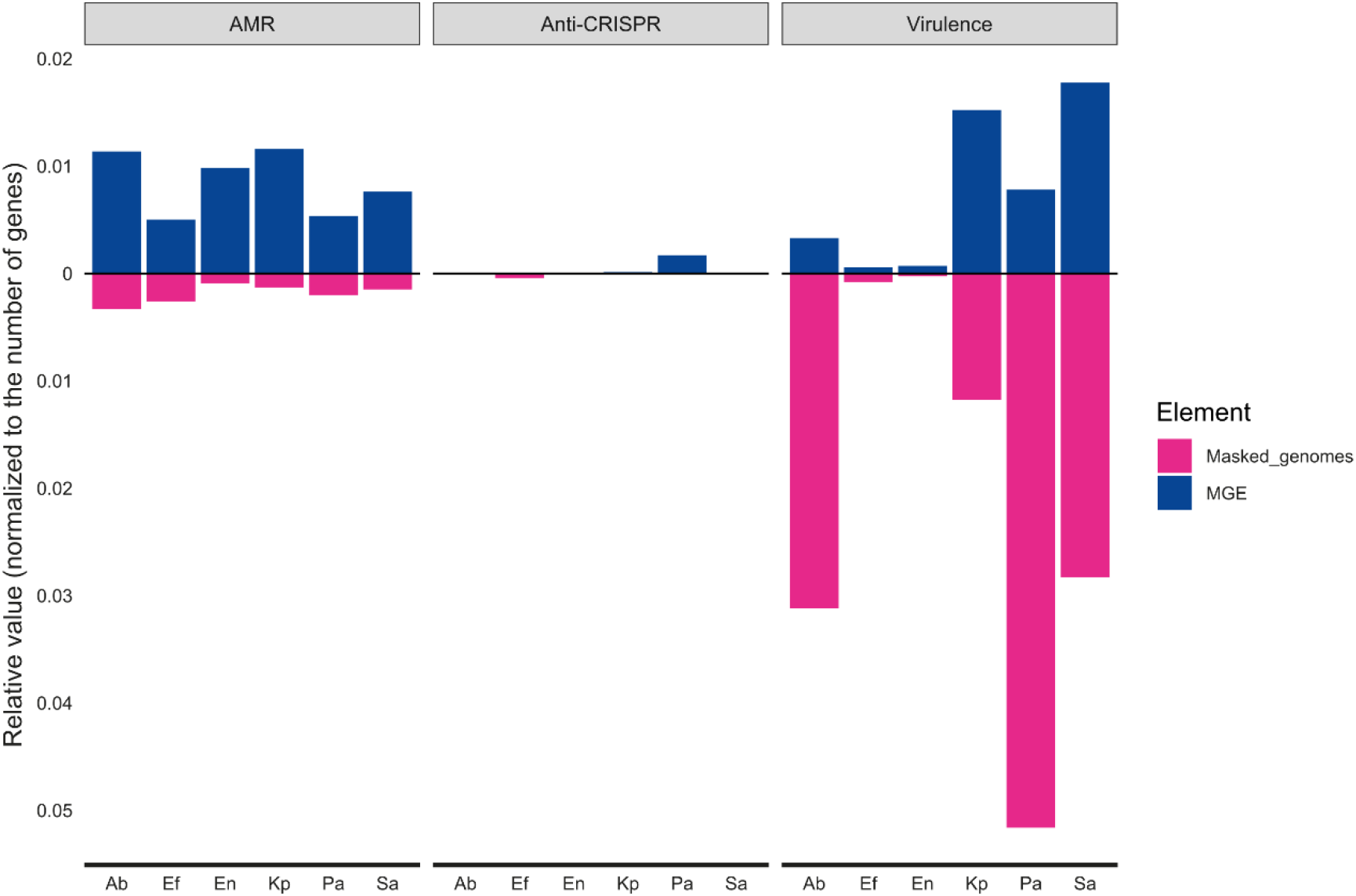
Relative proportion of AMR, virulence, and Anti-CRISPRs between MGEs and masked genomes across the ESKAPE pathogens. The number of AMR, virulence, or Anti-CRISPRs proteins found for MGEs or masked genomes per ESKAPE was normalized to the total number of proteins found for each element per ESKAPE pathogen.

### 5. CRISPR-Cas systems shape the number of MGEs, AMR, and virulence genes across ESKAPE pathogens

CRISPR-Cas systems were identified in every ESKAPE pathogen except *E. faecium*, which was therefore excluded from subsequent analyses. The proportion of genomes with CRISPR-Cas systems varied across the ESKAPE pathogens, from around 45.7% for *P. aeruginosa* to around 0.7% for *S. aureus* (**Figure 6A** and **Table S2**). We then explored the prevalence of CRISPR-Cas systems across closely related strains belonging to the same MLST profile. Since *Enterobacter* sp. consists of multiple species, this ESKAPE pathogen was excluded from subsequent analyses. Given the low prevalence of CRISPR-Cas systems in *S. aureus*, this species was also excluded, and we focused exclusively on *P. aeruginosa*, *K. pneumoniae*, and *A. baumannii*. Interestingly, some sequence types (ST) consisted entirely of either CRISPR-Cas positive or negative genomes (**Table S2**). For example, the most frequent MLST profile in *A. baumannii* from our dataset was ST2 (n=101), which only included strains with no CRISPR-Cas systems. On the other hand, the second most prevalent MLST profile in this species (ST1, n=14), only consisted of strains with I-F CRISPR-Cas systems. The most frequently observed MLST profile from *K. pneumoniae* (ST11, n=105), consists mostly of CRISPR-Cas negative strains (96.2%, 101/105). The four strains with positive hits carried IV-A3 CRISPR-Cas systems on plasmids. ST258 (n=47) was the second most common *K. pneumoniae* MLST profile identified in our dataset, and again, consisted entirely of strains with no CRISPR-Cas systems. However, ST147 and ST15 (n=31 and n=23, respectively) carried I-E CRISPR-Cas systems in all strains. Finally, looking at *P. aeruginosa*, we found that the most prevalent MLST profiles in our dataset (ST235 and ST549, n=16 and n=11, respectively) carried no CRISPR-Cas systems. The only exception was found in a ST235 strain, which carried an I-C CRISPR-Cas system on an ICE/IME. In contrast, ST233 and ST1971 (n=8 for both) consisted exclusively of strains carrying the I-F CRISPR-Cas system on masked genomes (**Table S2**). These findings suggest that the presence or absence of CRISPR-Cas systems across the ESKAPE pathogens is largely driven by the expansion of specific clones.

**Figure 6.**
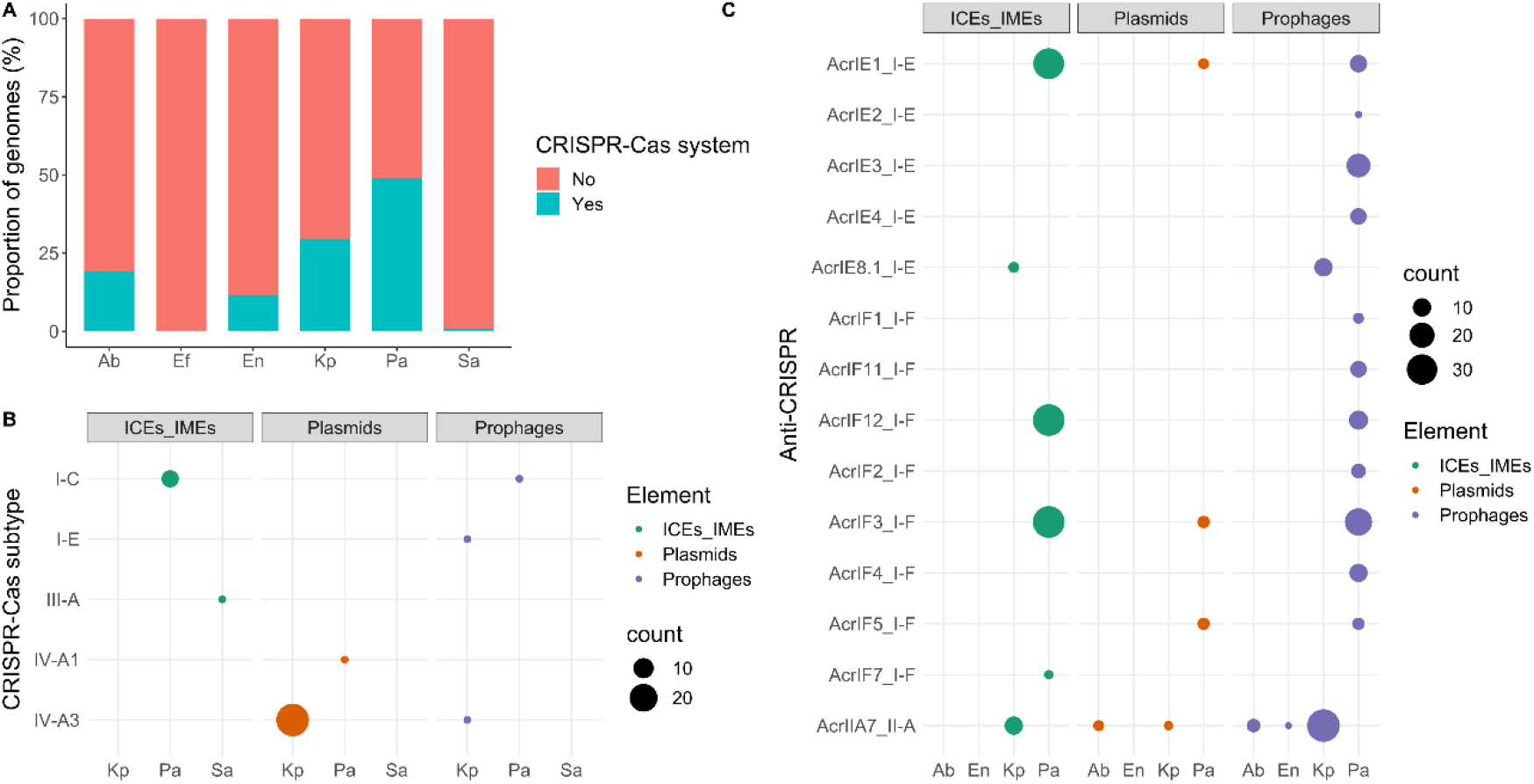
Distribution of CRISPR-Cas and Anti-CRISPR across the ESKAPE pathogens. **A)** Proportion of CRISPR-Cas positive genomes across the ESKAPE pathogens. **B)** Distribution of CRISPR-Cas systems within ICEs/IMEs, plasmids, and prophages. C) Distribution of Anti-CRISPR proteins across ICEs/IMEs, plasmids, and prophages. Even though we found Anti-CRISPRs on masked genomes, we only plotted those found on MGEs. A complete list of Anti-CRISPRs is given in Table S6. Ab, A. baumannii; Ef, E. faecium; En, Enterobacter sp.; Kp, K. pneumoniae; Pa, P. aeruginosa; Sa, S. aureus.

Our analysis revealed a large variety of MGE-encoded CRISPR-Cas subtypes, with I-C, I-E, I-F, III-A, IV-A1, and IV-A3 represented across the dataset (**Figure 6B**). We found CRISPR-Cas systems on plasmids (n=28 IV-A3 subtype in *K. pneumoniae* and n=1 IV-A1 in *P. aeruginosa*), ICEs/IMEs (n=7 I-C in *P. aeruginosa* and n=1 III-A in *S. aureus*), and prophages (n=1 I-C in *P. aeruginosa*, n=1 IV-A3 and n=1 I-E both in *K. pneumoniae*) (**Figure 6B** and **Table S6**). The plasmids and ICEs/IMEs carrying these systems were large, ranging from 102-430 kb. We also observed that all CRISPR-Cas-carrying plasmids have a MOB relaxase. This is in agreement with previous findings [55], which reported an enrichment of CRISPR-Cas systems across plasmids with conjugative functions and of larger sizes. We also found AMR and virulence genes on these CRISPR-Cas positive MGEs, but no anti-CRISPRs, suggesting that the CRISPR-Cas systems are functional.

When looking into the influence of GC content and sequence length in pairs of conspecific ESKAPE pathogens with and without CRISPR-Cas systems, we would expect to observe smaller and GC richer strains on those carrying these systems. Size expectations could only be met for *P. aeruginosa*, for which CRISPR-Cas positive genomes were significantly smaller than their counterparts (**Figure S12A**, p-value 0.0028), as observed before [56–58]. Surprisingly in *K. pneumoniae*, genomes with CRISPR-Cas systems were significantly larger than CRISPR-Cas negative genomes (**Figure S12A**, p-value 0.0023). The non-significant differences observed for *A. baumannii*, *Enterobacter* sp., and *S. aureus* could in part be explained by the low sample size of CRISPR-Cas positive genomes (**Figure 6A**). Regarding the GC content, we observed significant differences in *A. baumannii*, *K. pneumoniae*, and *P. aeruginosa*. CRISPR-Cas positive genomes were GC richer in *A. baumannii* and *P. aeruginosa* (**Figure S12B**, p-values 0.0013 and 0.046, respectively). Curiously, we noticed that CRISPR-Cas positive genomes in *K. pneumoniae* were GC poorer (**Figure S12B**, p-value 5.7e-09). Given the known association between GC content and genome size [47], these GC differences in CRISPR-Cas positive and negative *P. aeruginosa* genomes may be a spurious correlation driven by small size of CRISPR-Cas positive genomes. So, we corrected the GC content for variation in genome size, and we observed that the association was maintained (**Figure S13A**, p-value 0.0035), in accordance to a previous study [57].

Since virulence genes are overrepresented in the chromosome (**Figure 5**), we assessed the distribution of these genes in pairs of conspecific ESKAPE pathogens with and without CRISPR-Cas systems. Virulence genes were significantly less abundant in CRISPR-Cas positive genomes from *P. aeruginosa* and *A. baumannii* (**Figure S12C**, p-values 4.1e-06 and 0.0016, respectively). Given that *P. aeruginosa* genomes positive for these systems are significantly smaller than their CRISPR-Cas negative counterparts (**Figure S12A**), the lower prevalence of these genes in CRISPR-Cas positive *P. aeruginosa* genomes may again be driven by a spurious correlation. As so, we corrected the number of virulence genes for variation in genome size, and we observed that indeed the difference was no longer significant (**Figure S13B**, p-value 0.74), confirming our prediction that the genome size was a confounding variable obscuring the effect of CRISPR-Cas systems on the prevalence of virulence genes in *P. aeruginosa*.

We then explored whether CRISPR-Cas presence or absence reduced the number of MGEs acquired in pairs of conspecific ESKAPE pathogens. We would expect to detect a smaller number of MGEs in genomes harbouring these immune systems. However, this trend was only observed in *K. pneumoniae* (**Figure 7**). Finally, we focused on AMR and virulence genes carried exclusively by plasmids and ICEs/IMEs, as these were the most important vectors (**Figure S10**). For most MGE/ESKAPE pairs, we observed no significant difference between pairs of conspecific genomes with and without CRISPR-Cas systems. When it comes to AMR genes, we only observed significant differences in MGEs from *P. aeruginosa* (**Figure S14A**, p-values 0.037). Indeed, AMR genes were more prevalent on ICEs/IMEs from *P. aeruginosa* genomes with CRISPR-Cas systems (**Table S7**). A similar correlation was previously reported [58]. Curiously, the less prevalent I-C CRISPR-Cas subtype, which was exclusively identified in *P. aeruginosa* and mostly on ICEs/IMEs (**Table S6**), was recently found to be positively correlated with certain AMR genes [56]. Regarding virulence genes, we observed significant differences in MGEs from *K. pneumoniae*, where CRISPR-Cas-carrying elements were associated with more virulence genes than their CRISPR-Cas negative counterparts (**Figure S14B**, p-values 0.0054). Taken together, we observed species-specific trends shaping the number of MGEs, AMR, and virulence genes across pairs of conspecific ESKAPE genomes with and without CRISPR-Cas systems.

**Figure 7.**
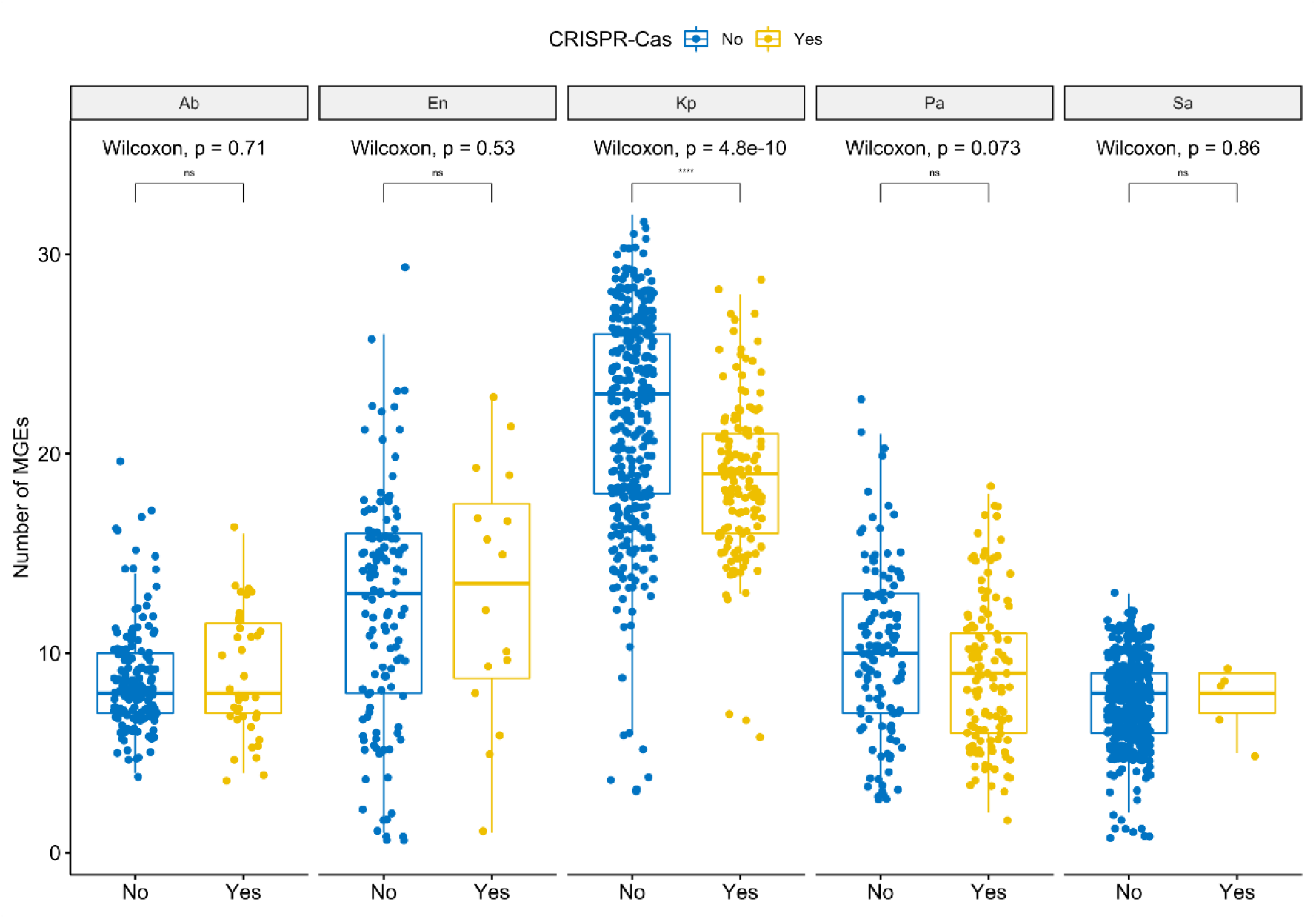
The absence of CRISPR-Cas systems does not associate with MGE increases in ESKAPE pathogens with the exception of *K. pneumoniae*. Boxplots compare the number of MGEs present in pairs of conspecific ESKAPE pathogens, with and without CRISPR-Cas systems. Values above 0.05 were considered as non-significant (ns). We used the following convention for symbols indicating statistical significance: * for p <= 0.05, ** for p <= 0.01, *** for p <= 0.001, and **** for p <= 0.0001. Ab, *A. baumannii*; En, *Enterobacter* sp.; Kp, *K. pneumoniae*; Pa, *P. aeruginosa*; Sa, *S. aureus*.

### 6. Anti-CRISPRs are overrepresented in the ESKAPE mobilome

Anti-CRISPR proteins (n=410) antagonising CRISPR-Cas subtypes I-E, I-F, and II-A were identified across prophages, ICEs/IMEs, and prophages from all ESKAPE pathogens except *S. aureus* (**Figure 6C** and **Table S8**). The majority of these counter-defence systems were found in prophages and ICEs/IMEs from *P. aeruginosa*. We also looked for these proteins across the masked genomes, and found hits in all ESKAPE except *A. baumannii* and *S. aureus* (**Table S8**). After correcting their prevalence to total number of genes, we verified that anti-CRISPRs are largely overrepresented in MGEs (∼15x) when compared with masked genomes (**Figure 5)**. When compared with masked genomes, our results show that Anti-CRISPR proteins are preferentially encoded in MGEs.

### 7. CRISPR spacers in ICEs/IMEs, prophages, and plasmids have different targeting biases

We explored the targets for all CRISPR spacers, retrieved from complete CRISPR-Cas systems, but also orphan CRISPRs, identified in our collection of ESKAPE genomes. Since we provided here a representative dataset of plasmids, ICEs/IMEs, and prophages, we used this collection as a database and took the CRISPR spacers identified in masked genomes as a query. In parallel, we used the CRISPR spacers identified in MGEs as a query and the MGEs collection as a database. For the latter, and to avoid self-targeting hits, we masked all CRISPR spacers from the MGEs collection used as database. We found a total of 1087 CRISPR spacers across the MGEs (n=554 on plasmids, n=343 on ICEs/IMEs, and n=190 on prophages. Only a small fraction of MGEs carried these spacers (1.3%, 33/2640 plasmids; 0.6%, 17/2685 ICEs/IMEs; and 0.07%, 11/16153, respectively). Given the large number of MGEs and CRISPR-Cas-encoding plasmids in *K. pneumoniae* (**Figure 1** and **Figure 6B**), more than half of the spacers were found in this species (577/1087). The large majority of plasmid spacers were identified on mobilizable plasmids (99.4%, 551/554).

We then looked for MGE spacer targets, and identified matches for 1271 MGEs from our collection (5.9%, n=1271/21478, **Table S9**). A substantially larger fraction of CRISPR spacers from plasmids targeted mobilizable plasmids from our ESKAPE collection (81.8%, 21628/26438 of total plasmid spacer’s interactions). Only a small fraction of plasmid spacers targeted prophages (13.0%), non-transmissible plasmids (4.7%), and ICEs/IMEs (0.5%) (**Figure 8A**). Most prophages spacers targeted other prophages (85.3%, 1513/1773 of total prophage spacer’s interactions). Only a small fraction of prophage spacers targeted ICEs/IMEs (7.6%) and plasmids (7.1%). Surprisingly and unlike CRISPR spacers found on plasmids and prophages, ICEs/IMEs spacers were not biased towards other ICEs/IMEs (37.8%, 795/2102 of total ICEs/IMEs interactions), but towards prophages (61.3%). Only a small fraction of ICEs/IMEs spacers targeted plasmids (0.9%) (**Figure 8A**).

**Figure 8.**
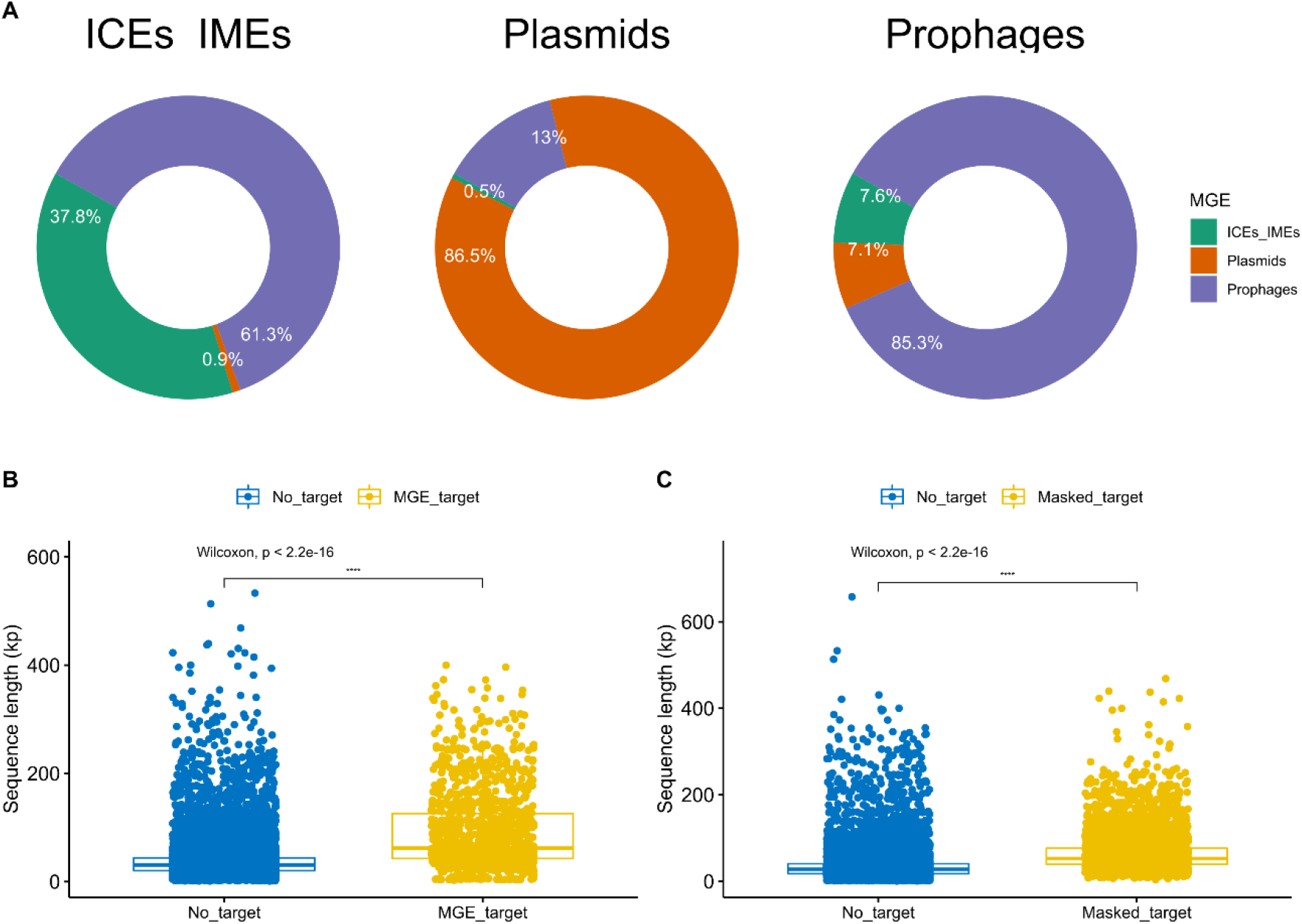
CRISPR spacers are involved in MGE-MGE conflict. **A)** CRISPR spacers found on ICEs/IMEs (top left), plasmids (top middle), and prophages (top right), and their interactions with spacer targets identified in ESKAPE MGEs. **B)** Boxplots comparing the sequence length of MGEs targeted by CRISPR spacers found in MGEs. **C)** Boxplots comparing the sequence length of MGEs targeted by CRISPR spacers found in masked genomes. Values above 0.05 were considered as non-significant (ns). We used the following convention for symbols indicating statistical significance: * for p <= 0.05, ** for p <= 0.01, *** for p <= 0.001, and **** for p <= 0.0001. Ab, *A. baumannii*; En, *Enterobacter* sp.; Kp, *K. pneumoniae*; Pa, *P. aeruginosa*; Sa, *S. aureus*.

When it comes to CRISPR arrays on masked genomes, we observed a total of 13400 spacers in 30.4% ESKAPE genomes (n=531/1746). Even though the prevalence of CRISPR-Cas systems in *S. aureus* was considerably low (< 1%, **Figure 6A**), we found spacers in 30.7% of the total *S. aureus* strains (n=170/553). Consistent with the absence of CRISPR-Cas systems in the *E. faecium* strains from our dataset, no spacers were identified in masked genomes from this species. The number of CRISPR spacers per genome varied considerably between the masked genomes of the ESKAPE pathogens (**Figure S15**), reaching as high as 189 in *A. baumannii*. In fact, only strains from this species carried more than 100 spacers per masked genome. We observed that 38.4% (5141/13400) of CRISPR spacers in ESKAPE masked genomes yielded matches to spacer targets in MGE sequences, found on 16.4% MGEs (n=3523/21478, **Table S10**). Most CRISPR spacers from masked genomes targeted prophages (44.5%, 83034/186619 total interactions), mobilizable plasmids (39.9%), and ICEs/IMEs (10.5%). As observed for CRISPR spacers in MGEs, CRISPR spacers from masked genomes rarely targeted non-transmissible plasmids (5.2%). We then tested if MGE or masked genome spacers preferentially targeted ESKAPE MGEs of variable size. We found that the CRISPR spacers from both MGEs and masked genomes targeted significantly larger MGEs than those with no spacer targets (p-value <2.2e-16**, Figures 8B and 8C**). Altogether, our results show that plasmids and prophages mostly targeted other plasmids and prophages, respectively, while ICEs/IMEs preferentially targeted prophages. Our data also shows that CRISPR spacers found either on MGEs or masked genomes consistently target larger MGEs.

## Discussion

In this work, we performed a systematic analysis of prophages, ICEs/IMEs, and plasmids across all ESKAPE pathogens. We focused on this panel because the ESKAPE group of pathogens consists of clinically-relevant bacteria, for which many genomes are completely sequenced (an important parameter when delineating MGEs), and which include representatives of both Proteobacteria and Firmicutes, and also phylogenetically divergent bacteria (with exception of *K. pneumoniae* and *Enterobacter* sp., the remaining representatives belong to different bacterial families). Studying MGEs in parallel allowed us to explore their uneven distribution across a collection of complete genomes from important pathogens, and to explore potential DNA sharing events between different MGE types. By separating these elements from masked (MGE-free) genomes, we were able to observe an overrepresentation of AMR genes and anti-CRISPRs across the ESKAPE mobilome. Furthermore, we assessed the influence of CRISPR-Cas defence systems in shaping the acquisition of MGEs and beneficial genes, and we unveiled different targeting biases for CRISPR spacers identified in plasmids, prophages, and ICEs/IMEs.

The network-based approach used here to study the ESKAPE MGEs revealed a clear structural differentiation, where the majority of clusters were homogeneous for a given ESKAPE/MGE pair. Using pairwise genetic distances of alignment-free *k*-mer sequences has circumvented the exclusion of non-coding elements that was observed in gene content similarity networks from previous work [59], providing a more comprehensive picture of plasmid population and dynamics [13]. Other groups have shown that plasmids form coherent clusters [13,14], similar in concept to what was observed for bacterial genomes [48,49,60]. Focusing on the ESKAPE pathogens, we demonstrated here that the same happens for ICEs/IMEs and prophages for the majority of the clusters. However, two large heterogeneous clusters were also observed. Unlike bacterial genomes, where recombination between closely related replicons is the main force promoting genomic cohesiveness [48,60] (**Figure S6**), MGEs such as plasmids, ICEs/IMEs and prophages are capable of exchanging DNA beyond the genus barrier (and even occasionally across phylum) [61]. Based on our network-based approach, we observed that the ESKAPE mobilome appears to follow a bipartite structure, with some elements being capable of shuffling DNA between distantly related species.

The abundance of MGEs is strongly associated with the prevalence of AMR genes. This is particularly evident for *K. pneumoniae*, which carries a high proportion of important vectors of AMR genes, such as plasmids and ICEs/IMEs. After correcting the prevalence to total number of genes, we observed that AMR genes are nearly 5 times more likely to be found on ESKAPE MGEs than on masked genomes. Most likely the use of different antibiotics targeting either Gram negative or positive infections may have selected for the emergence of different AMR genes across the Proteobacteria/Firmicutes divide. For example, vancomycin is used as last resort for the treatment of sepsis and other infections caused by Gram-positive bacteria, while colistin mainly serves to target multi-drug resistant Gram-negative infections [30]. Anti-CRISPRs are nearly 15 times more abundant on the ESKAPE MGEs than on masked genomes. Unlike AMR genes, where genes conferring resistance to similar antibiotics were identified in different ESKAPE pathogens, the distribution of virulence genes across the ESKAPE mobilome was mostly species-specific. This may be explained by different mechanisms of virulence and toxicity across bacterial species. Even considering the relative proportion of these genes, we confirmed that these genes are twice more likely to be identified in masked genomes than in MGEs.

Curiously, we found no CRISPR-Cas systems in our curated *E. faecium* dataset. In addition to our main analysis, we specifically searched for the presence of these systems on the excluded genomes, based on <95% genome similarity threshold defined for species delineation [60], and found three strains with these systems (**Table S15**). Since the majority of the most representative MLST profiles in our collection consists of genomes either with or without CRISPR-Cas systems (**Table S2**), analysis of intra-ST CRISPR variability between pairs of conspecific ESKAPE genomes was not performed in this study. Except for *P. aeruginosa* [56,57], we found no evidence for genome length as a marker for HGT inhibition by CRISPR-Cas systems. A similar observation was made before, for a different collection of bacterial pathogens [58].

Defence systems such as CRISPR-Cas are inherently costly to bacterial hosts, mainly due to different forms of autoimmunity [62]. To offset the short-term deleterious effects, these systems benefit from associating with MGEs, such as the examples observed here and elsewhere [41]. Although these and other defence mechanisms such as restriction-modification systems do not qualify as bona fide MGEs (the systems lack mechanisms controlling their own replication), these quasi-autonomous systems take advantage of MGEs to promote their own dissemination and maintenance across bacterial hosts. Conversely, MGEs benefit from these systems and may pervasively repurpose them for inter-MGE competition [55]. In fact, we found that spacers in the ICE/IME, prophage, and plasmid CRISPR arrays target competing MGEs, underscoring the genetic independence of CRISPR-Cas systems in MGE-MGE conflicts. Importantly, the presence of CRISPR-Cas subtypes preferentially distributed in MGEs (I-C and IV-A3) points to the existence of distinct selective pressure that promote the maintenance of specific subtypes on ICEs/IMEs and plasmids versus masked genomes. Given the large sizes of CRISPR-Cas systems, we observed a bias in their distribution towards larger MGEs (both plasmids and ICEs/IMEs >100kb). Since plasmids with an identifiable relaxase (hence classified as conjugative or mobilizable) are larger than the so-called non-transmissible plasmids [44], we unsurprisingly found a relaxase in all CRISPR-Cas positive plasmids. Prophages seem to follow similar streamlining dynamics [63]. Even though nearly half of *P. aeruginosa* genomes carry at least one CRISPR-Cas system (**Figure 6A** and **Table S2**), multiple anti-CRISPRs were found across prophages and ICEs/IMEs (**Figure 6C**), suggesting a potential role played by these elements in silencing immune systems in this species.

Our results shows that only a restricted fraction of CRISPR spacers matched spacer MGE targets, which is in agreement with previous findings [55,57]. While the large majority of plasmid- and prophage-encoded spacers were predicted to target other plasmids and prophages, respectively, CRISPR arrays in ICEs/IMEs preferentially targeted prophages, but also a large proportion of other ICEs/IMEs (**Figure 8**). This complementary targeting preference can be explained by the different lifestyle of these MGEs. Since plasmids are maintained as extra-chromosomal elements, and ICEs/IMEs and prophages are integrated in the chromosome, we hypothesize that while plasmids preferentially target plasmid competitors [41], ICEs/IMEs exploit CRISPR-Cas systems to protect their host against viral predators and other ICEs/IMEs.

Several bioinformatic tools exist to look for plasmids and prophages, but currently the options for ICEs/IMEs are scarce [64]. We provide here an accurate identification of these elements, building upon a recently reported tool to scan RGPs [65]. However, our approach depends on the availability of a pangenome for the considered taxa, which is an important limitation for species with an insufficient number of completely sequenced genomes. Moreover, our work only focused on three MGE types, which are likely to be of main importance for the dissemination of genes involved in pathogenesis and AMR and thus critical for our understanding of the evolution of the ESKAPE pathogens. Nevertheless, other MGEs should be considered in future work, for example those that contribute to communication between cells (such as phage-inducible chromosomal islands [66]), or intracellular transfer events (e.g., transposons and insertions sequences [19]).

To conclude, our results indicate that prophages, ICEs/IMEs, and plasmids are asymmetrically distributed across the ESKAPE pathogens. We found that these MGEs can exchange DNA beyond the genus (and phylum) barrier, even though most clusters are constrained by host similarity and/or the type of MGE. We observed that the proteome of ESKAPE MGEs is highly diverse, involved in diverse functional categories, and features convoluted distribution patterns (including both MGE/ESKAPE specific and widespread proteins). When comparing against masked (MGE-free) genomes, we observed the pervasive association of AMR genes and anti-CRISPRs with the ESKAPE mobilome. We also found different targeting biases to CRISPR spacers found on plasmids, prophages, and ICEs/IMEs, highlighting their genetic independence. Taken together, our results illustrate the potential of network-based approaches and comparative genomics to underscore the composition and dynamics of gene flow across different MGEs, and sheds a new light in the role of the overlooked ICEs/IMEs as important players in the MGE-MGE warfare in the ESKAPE pathogens and thus the main groups of highly problematic human pathogens.

## Material and methods

### ESKAPE pathogens collection

We retrieved all complete ESKAPE genomes available in the NCBI Reference Sequence Database (RefSeq, accessed on 12-11-2020), using ncbi-genome-download v0.3.0 (https://github.com/kblin/ncbi-genome-download). Genomes listed as ‘unverified’ were removed from our dataset. We also excluded genomes classified as ‘Enterobacteriaeceae’. Finally, we used pyANI v0.2.10 (https://github.com/widdowquinn/pyani) to calculate the average nucleotide identity based on MUMmer (ANIm), and removed genomes with an ANIm value below the 95% threshold for species delineation [60,67]. To evaluate the taxonomy of the *Enterobacter* species, we retrieved genomes for *Enterobacteriaceae* type strains and used them together with the *Enterobacter* genomes to create a phylogenetic tree using GToTree v1.5.22 (https://github.com/AstrobioMike/GToTree) [68] and the IQ-TREE algorithm to estimate maximum likelihood [69]. We used the pre-built set of 74 single copy gene bacterial Hidden Markov Models (HMM) available in GToTree. Genomes labelled as belonging to the *Enterobacter* genus, but clustered in the phylogenomic tree with type strains other than those from the *Enterobacter* genus, were removed from subsequent analyses. MLST profiles were determined with mlst v2.19.0 (https://github.com/tseemann/mlst). The curated genomes were automatically annotated using Prokka v1.14.6 (https://github.com/tseemann/prokka) [70].

### Extraction of plasmids, ICEs/IMEs, and prophages

Since the ESKAPE pathogens (as most bacteria) are haploid, we separated the large replicon (i.e., the chromosome) from the extrachromosomal replicons. For the latter, only accessions with ‘plasmid’ and ‘complete sequence’ on their description were kept and were used for further plasmid analyses.

To extract ICEs from chromosomal replicons, we used the chromosomal genbank files created with Prokka as input to build a pangenome for each ESKAPE pathogen. We used ppanggolin v1.1.96 (https://github.com/labgem/PPanGGOLiN)[71], which uses a graphical model and a statistical method to partition the pangenome in persistent, shell and cloud genomes. We used the panRGP method to build the pangenomes [65]. This method predicts RGPs, which are clusters of genes made of accessory genes (shell and cloud genomes) in the pangenome graph. We then used bedtools v2.30.0 (https://bedtools.readthedocs.io/en/latest/) [72] to extract RGPs from the chromosomal replicons. The proteomes of these extracted RGPs were scanned for relaxases with hmmer v.3.3.1 (http://hmmer.org/) [73] against MOBfamDB, a curated relaxase profile HMM database [74]. Simultaneously, the proteomes were screened with hmmer against integrases (Pfam accession PF00589) and recombinases (PF07508). RGPs with hits both for integrases (phage integrase or recombinase) and relaxases were classified as putative ICEs/IMEs and were kept for further analysis.

To look for prophages, we masked the ICEs/IMEs locations in the chromosomal replicons using bedtools. The ICE/IME-masked chromosomes were annotated with Prokka, using as proteins of interest a collection of non-redundant viral proteins downloaded from NCBI’s RefSeq database (https://ftp.ncbi.nlm.nih.gov/refseq/release/viral/, accessed on the 25/01/2022). The masked chromosomal genbank files were then used as input in phispy v4.2.6 (https://github.com/linsalrob/PhiSpy), which combines similarity- and composition-based strategies to look for prophages [75]. We also masked the prophage regions in the chromosomal replicons, to build the final masked genomes, that are ICE/IME- and prophage-masked (these masked replicons are also free of plasmids, since these are part of the extrachromosomal replicons).

### Network-based approach

To estimate the pairwise distances between all ESKAPE MGEs (plasmids, ICEs and prophages), we first ran the MMSEQseqs2 v13.45111 package [76], using 90% sequence identity for clustering each MGE type. We then reduced the dereplicated MGEs into sketches and compared the Jaccard index (JI) and mutation distances between pairs of MGEs using BinDash v 0.2.1 (https://github.com/zhaoxiaofei/bindash)[77]. Each MGE sequence was converted to a set of 21-bp *k*-mers. We used the mean() and median() functions in R to calculate the arithmetic mean and median of the JI, respectively. Only JI equal to or above the mean and median were considered, and the mutation distances were used as edge attributes to plot the network with Cytoscape v3.9.0 under the prefuse force directed layout (https://cytoscape.org/). We used the Analyzer function in Cytoscape to compute a comprehensive set of topological parameters, such as the clustering coefficient, the network density, the centralization, and the heterogeneity.

### Functional annotation

COGs annotation of the MGE proteins was carried out through sequence alignments against the COGs 2020 database (https://www.ncbi.nlm.nih.gov/research/cog-project/). The alignments were performed with DIAMOND v0.9.10.111 [78] with a cutoff evalue of 1e-05 and 80% coverage of both query and subject sequences. The COGs database was set up using a python script (https://github.com/kkpenn/merger_COG2020/blob/main/merger_2.py) and DIAMOND makedb with default settings. Around 36, 38 and 55% of the proteins encoded in plasmids, ICEs/IMEs, and prophages matched a protein in the COGs database, respectively, and were therefore annotated with the information of their corresponding homolog. Lists of non-redundant COG definitions (e.g. COG0105) were extracted separately for prophages, plasmids and ICEs/ IMEs, and compared with venny v2.1.0 (https://bioinfogp.cnb.csic.es/tools/venny/index.html) to identify unique and shared COGs. Likewise, COGs occurrence was determined separately for the proteomes of the three MGE types in the different ESKAPE. Information on the COGs classification into functional categories was retrieved from https://ftp.ncbi.nih.gov/pub/COG/COG2020/data/. The relative frequency of the different COG functional categories per MGE/ESKAPE pair was calculated by summing up the occurrences of COGs belonging to a given functional category and dividing the resulting number by the total number of proteins observed in the corresponding MGE/ESKAPE pair.

To explore the diversity of MGE-encoded proteins, we combined their proteomes (943246 proteins) and clustered them using the cluster algorithm from the MMseqs2 v13.45111 package [76]. The proteins were clustered at 80% sequence identity, 80% coverage, and otherwise default settings to match the parameters used by ppanggolin when generating the ESKAPE pangenomes. The relative frequency of the different protein clusters per MGE/ESKAPE pair was calculated following the same approach used to estimate the relative frequency of COG functional categories but using the occurrence of proteins belonging to a given cluster instead. Representatives of the 72247 clusters identified were annotated with eggNOG-mapper v2 [79] with default settings to explore the functions of the MGE-encoded proteins further.

We used abricate v1.0.1 (https://github.com/tseemann/abricate) to scan extracted MGEs and masked genomes against antimicrobial resistance and virulence genes (using pre-downloaded databases from Resfinder[80] and VFDB[81] containing 3138 and 4329 sequences, respectively, and both updated on the 28-03-2022). To identify and classify CRISPR-Cas systems, we used CRISPRCasTyper v1.2.3 (https://github.com/Russel88/CRISPRCasTyper)[82]. We also used this tool to look for CRISPR spacers. The entire CRISPR arrays identified on MGEs were then masked using bedtools, and these masked MGEs served as a local blast database using blast v2.12.0 (https://blast.ncbi.nlm.nih.gov/Blast.cgi?PAGE_TYPE=BlastDocs), when using MGE CRISPR spacers as a query. CRISPR spacers from masked genomes were also blasted against a local database of our extracted (non-masked) MGEs. Hits with at least 95% nucleotide identity and 95% sequence coverage were considered as spacer targets [83]. While a representative collection of plasmids and virus is publicly available at RefSeq’s NCBI database (n=33269 and 13778, respectively, accessed on the 21/05/2021), a substantially smaller collection of ICEs/IMEs is available at ICEberg database (n=1325), and was last updated in September 2018. Due to this limitation in the number of publicly available ICEs/IMEs sequences, and to have a representative collection of these three different types of MGEs, we focused on the curated dataset presented in this study to look for targets of CRISPR spacers.

We retrieved an Anti-CRISPR collection of 1111 non-redundant proteins from Anti-CRISPRdb v2.2 (http://guolab.whu.edu.cn/anti-CRISPRdb/, accessed on the 29-03-2022). This collection was used to build a local database with diamond v2.0.14 (https://github.com/bbuchfink/diamond) [78]. We used the blastp command in diamond to scan the MGEs and masked proteomes against the anti-CRISPR local database. We used an amino acid-based homology approach to find Anti-CRISPRs encoded in the ESKAPE mobilome. Even though recent approaches have applied a guilt-by-association method to identify new Anti-CRISPRs [29], currently there is no tool available to apply this method in a large dataset of bacterial genomes.

### Statistical analysis

Comparisons between MGEs’ GC content and sequence length were performed using the Kruskal-Wallis test, and the p-values adjusted using the Holm–Bonferroni method. Comparisons between pairs of conspecific genomes with and without CRISPR-Cas systems, as well as between MGE targets for CRISPR spacers, were performed using the Wilcoxon test, and the p-values adjusted using the Holm–Bonferroni method. Values above 0.05 were considered as non-significant (ns). We used the following convention for symbols indicating statistical significance: * for p <= 0.05, ** for p <= 0.01, *** for p <= 0.001, and **** for p <= 0.0001.

### Code availability

Analyses were made with a combination of shell and R 4.0.3 scripting. R scripts and supporting tables used to create the figures are available at https://gitlab.gwdg.de/botelho/eskape_paper).

## Acknowledgments

We would like to thank José Penades for comments and suggestions. We are grateful for funding from the Max-Planck Society and the Max-Planck Institute for Evolutionary Biology in Ploen (Fellowship to HS), the German Science Foundation (funding under Germany’s Excellence Strategy EXC 2167-390884018 as well as the Research Training Group 2501 TransEvo to HS), and the Leibniz ScienceCampus Evolutionary Medicine of the Lung (EvoLUNG, to HS). This work was supported by the Kiel Life Science Postdoc Award to JB. This research was supported in part through high-performance computing resources available at the Kiel University Computing Centre.

---

**Figure S1.**
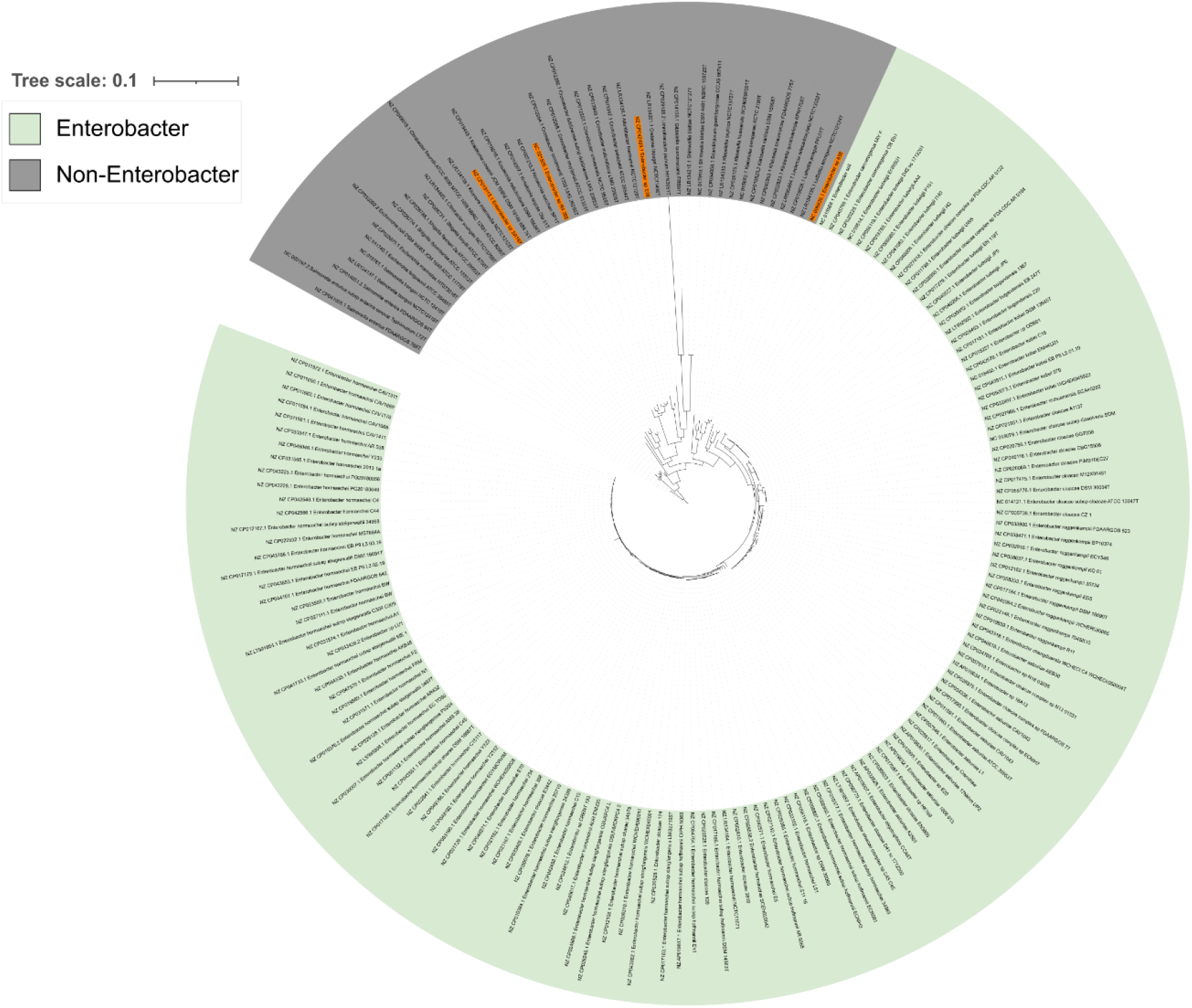
Maximum likelihood tree representing type strains belonging to the *Enterobacteriaceae* family and all complete genomes classified as *Enterobacter*. Genomes that are positioned in the non-*Enterobacter* cluster (gray), and poorly classified as belonging to the *Enterobacter* genus, were highlighted in orange.

**Figure S2.**
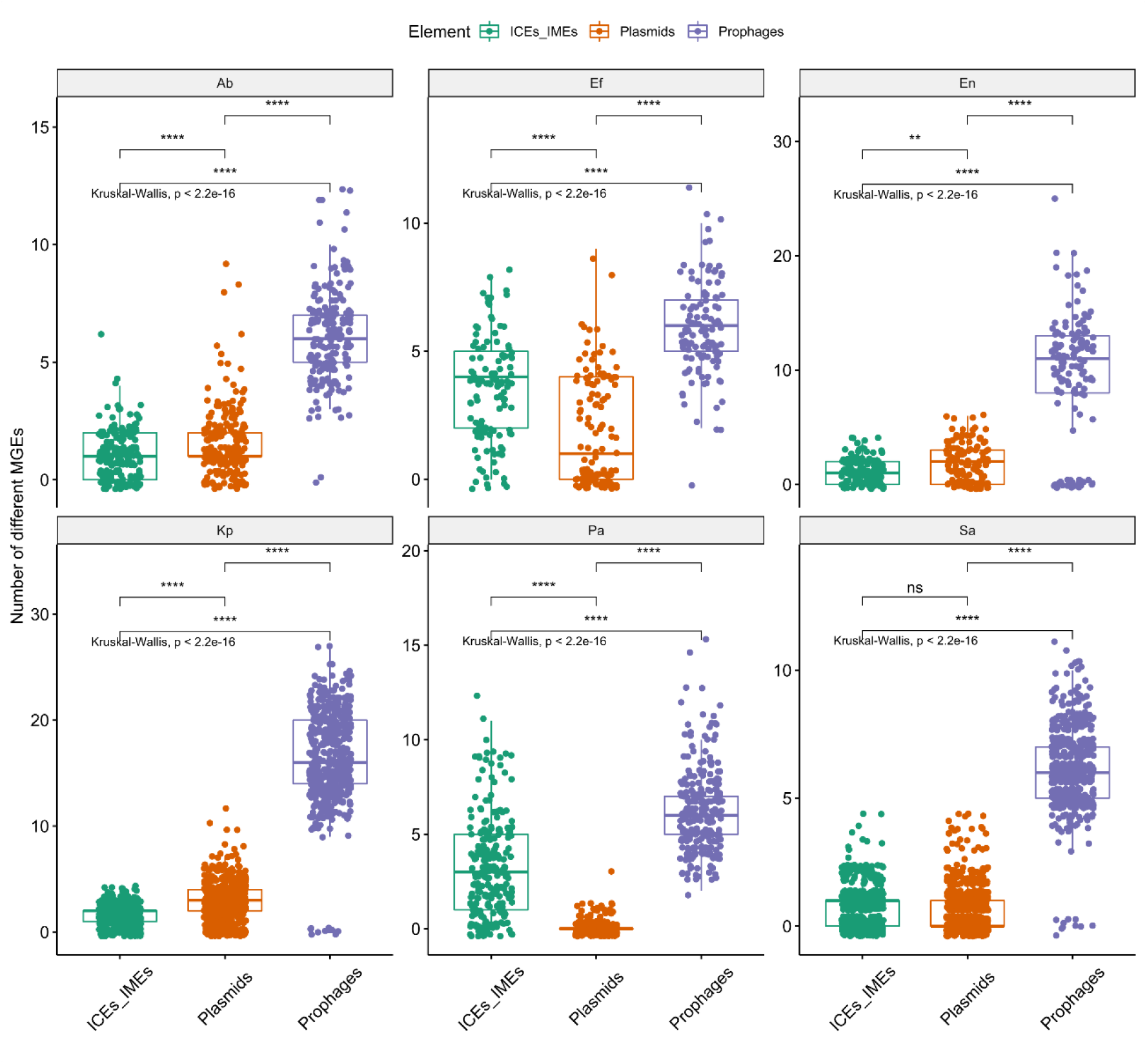
Boxplots representing the number of prophages, ICEs/IMEs, and plasmids per genome across the ESKAPE pathogens. Ab, *A. baumannii*; Ef, *E. faecium*; En, *Enterobacter* sp.; Kp, *K. pneumoniae*; Pa, *P. aeruginosa*; Sa, *S. aureus*. Values above 0.05 were considered as non-significant (ns). We used the following convention for symbols indicating statistical significance: * for p <= 0.05, ** for p <= 0.01, *** for p <= 0.001, and **** for p <= 0.0001.

**Figure S3.**
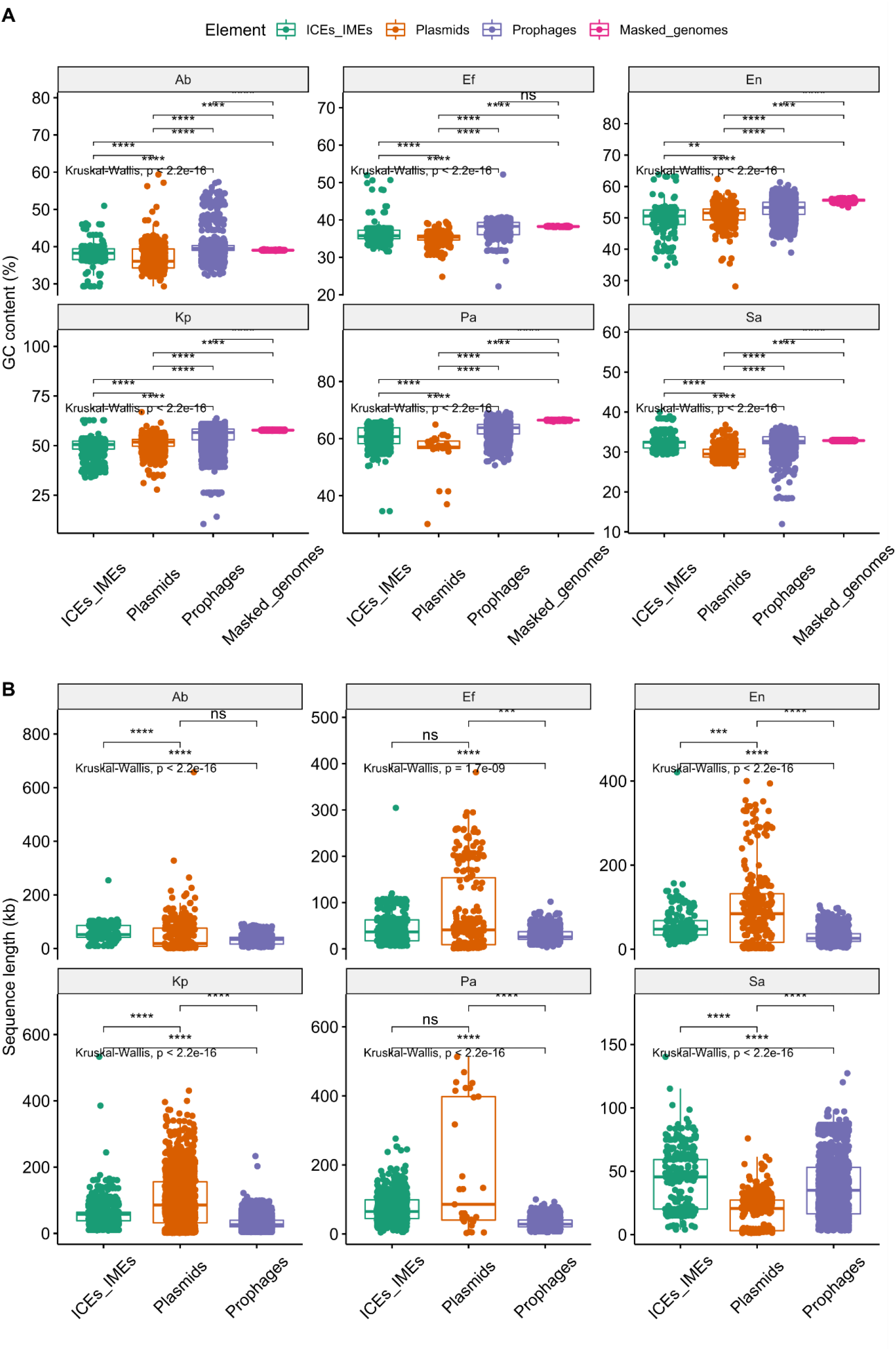
Variation in **A)** GC content and **B)** sequence length between MGEs and masked genomes. Ab, *A. baumannii*; Ef, *E. faecium*; En, *Enterobacter* sp.; Kp, *K. pneumoniae*; Pa, *P. aeruginosa*; Sa, *S. aureus*. Values above 0.05 were considered as non-significant (ns). We used the following convention for symbols indicating statistical significance: * for p <= 0.05, ** for p <= 0.01, *** for p <= 0.001, and **** for p <= 0.0001.

**Figure S4.**
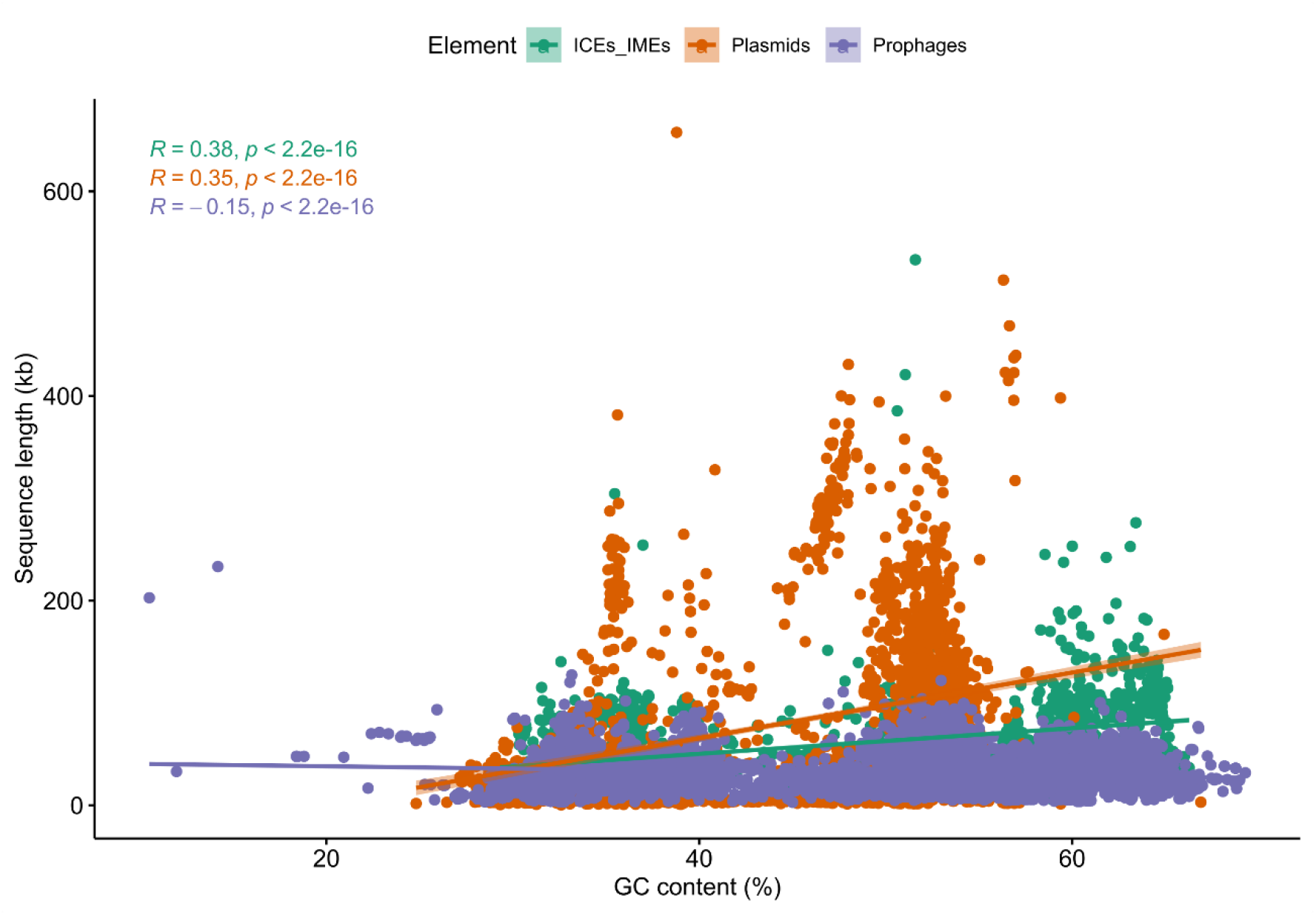
Scatter plot between the different MGEs’ sequence length and GC content. Regression lines for each MGE type are shown in different colours. Confidence interval is displayed around smooth. Level of confidence interval is 0.95.

**Figure S5.**
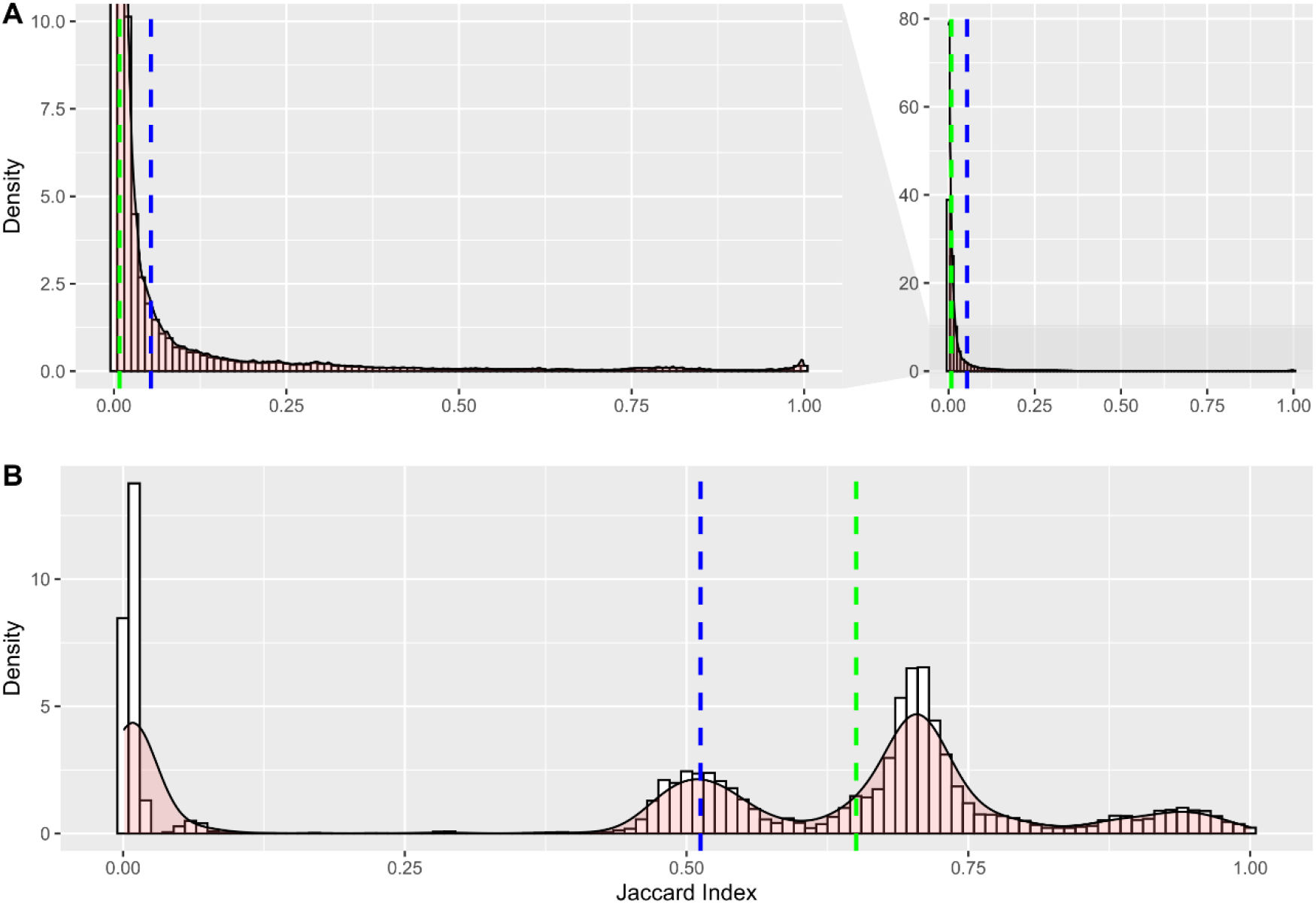
Density plot showing the distribution of Jaccard index (JI) values for the **A)** MGE and **B)** masked genomes network. The mean distribution for the JI is shown in blue, while the median is shown in green. In **A)**, density values from 0-10 were zoomed in and shown in the left.

**Figure S6.**
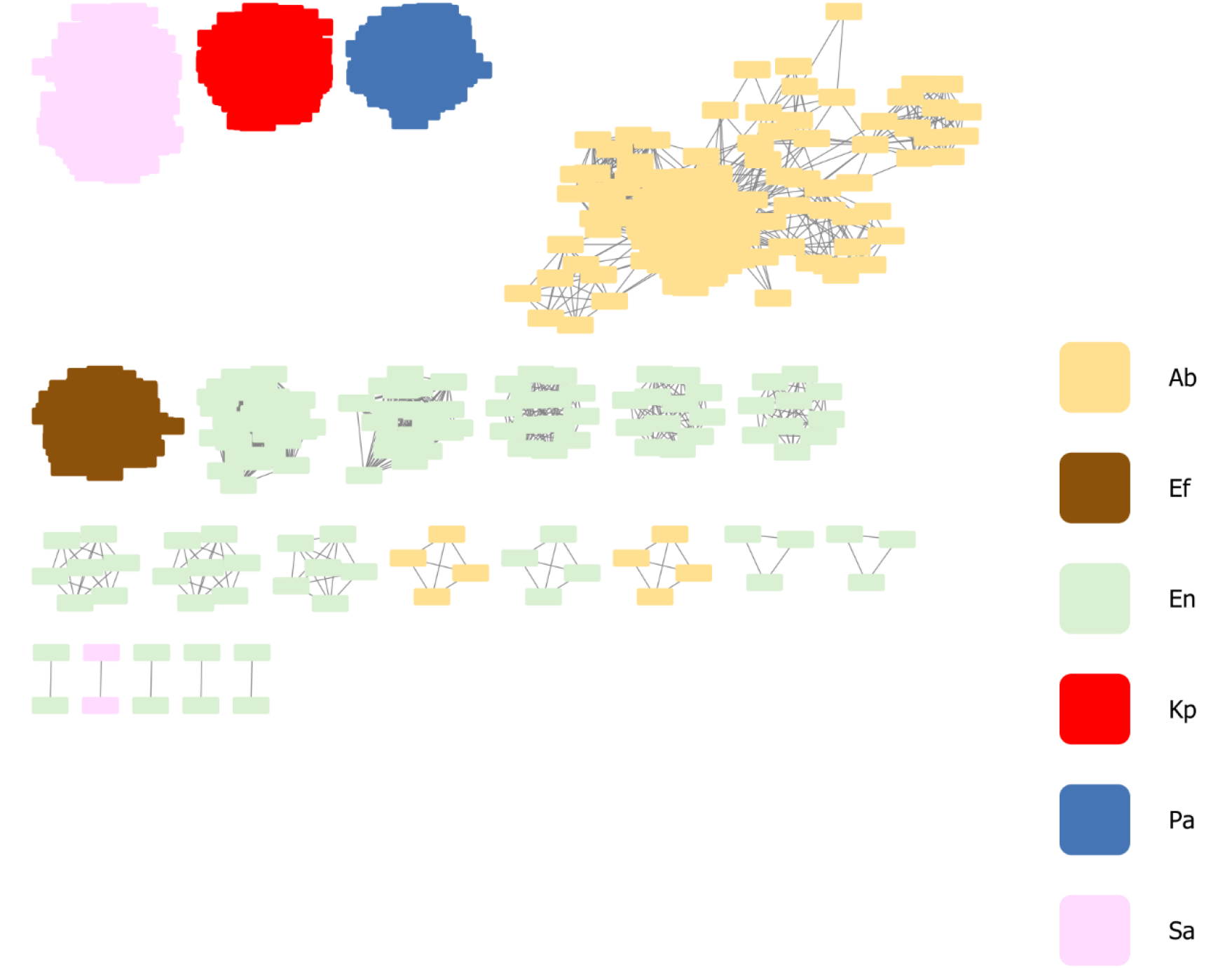
Network of clustered masked genomes grouped by ESKAPE pathogen, using the mean Jaccard index as a threshold. Each masked genome is represented by a node, connected by edges according to the pairwise distances between all masked genome pairs. Ab, *A. baumannii*; Ef, *E. faecium*; En, *Enterobacter* sp.; Kp, *K. pneumoniae*; Pa, *P. aeruginosa*; Sa, *S. aureus*.

**Figure S7.**
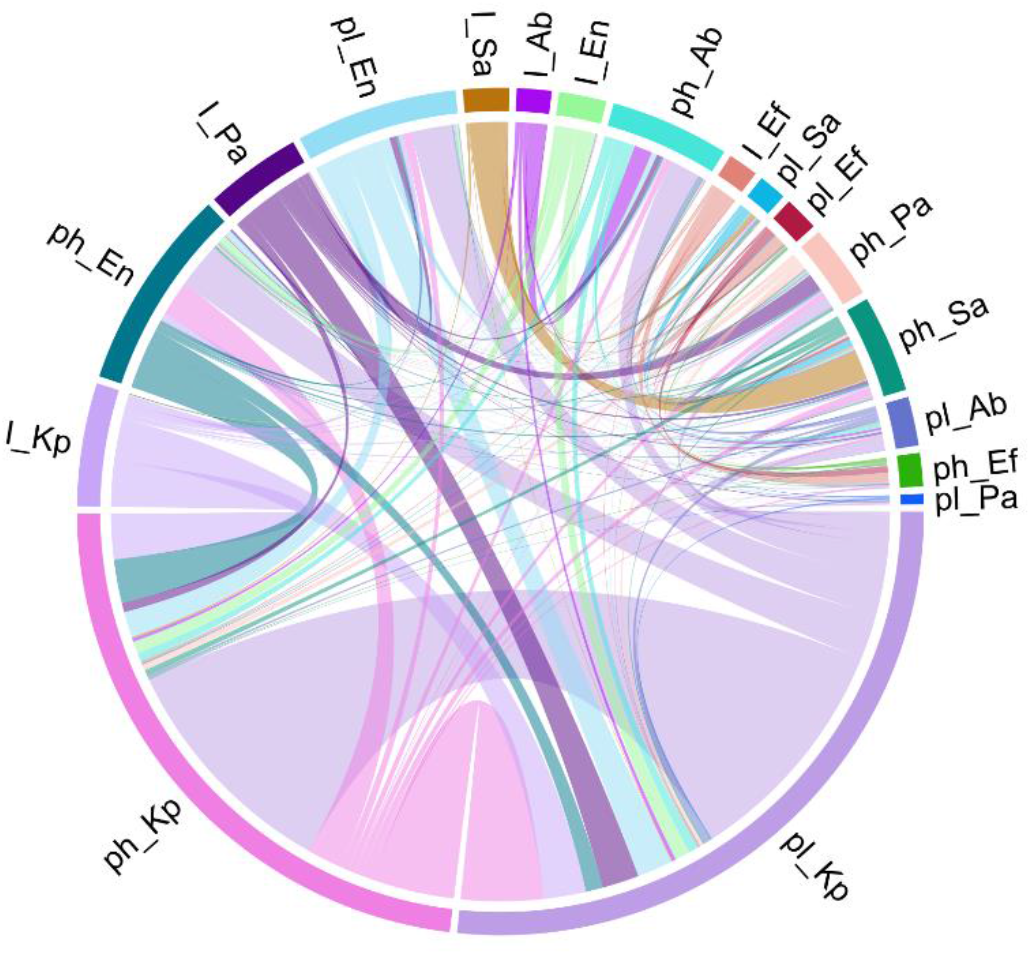
Chord diagram showing DNA sharing events between different ESKAPE pathogens. The width of sectors represents total number of inferred interactions between two different MGE/ESKAPE pairs. The width of links are proportional to the number of inferred DNA sharing events. Interactions between the same ESKAPE/MGE pairs were excluded. Ab, *A. baumannii*; Ef, *E. faecium*; En, *Enterobacter* sp.; Kp, *K. pneumoniae*; Pa, *P. aeruginosa*; Sa, *S. aureus*.

**Figure S8.**
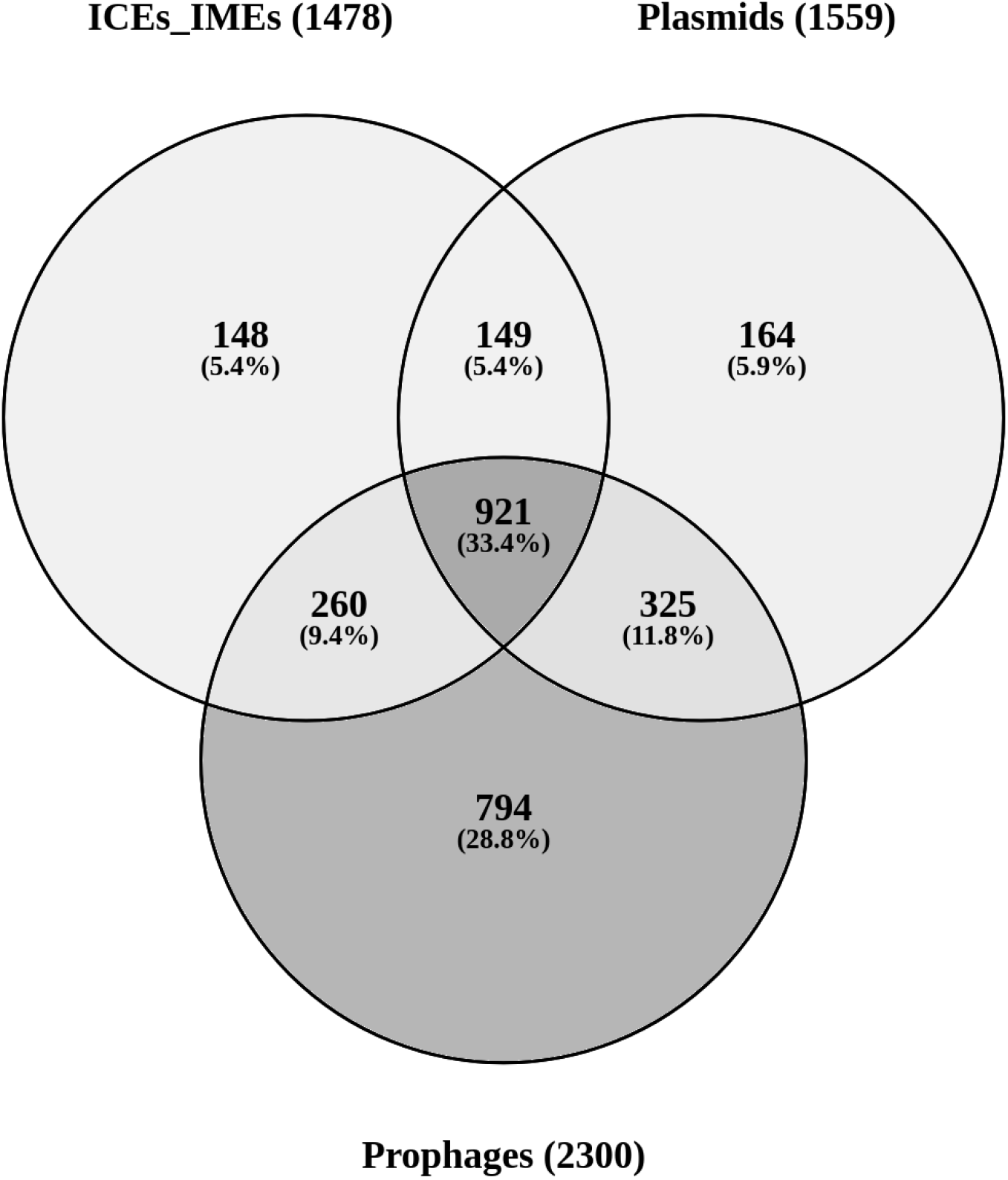
COGs identified in ESKAPE MGEs. Venn diagram illustrating the distribution of 2761 different COGs identified in ESKAPE ICEs/IMEs, plasmids and phages. The number of COGs detected for each MGE type is indicated in parenthesis. Note that the numbers displayed in the figure do not correspond to the total number of proteins with an identifiable COG in the MGE sequences analysed but to the distinct COG definitions (e.g. COG0105) detected in the MGE proteomes.

**Figure S9.**
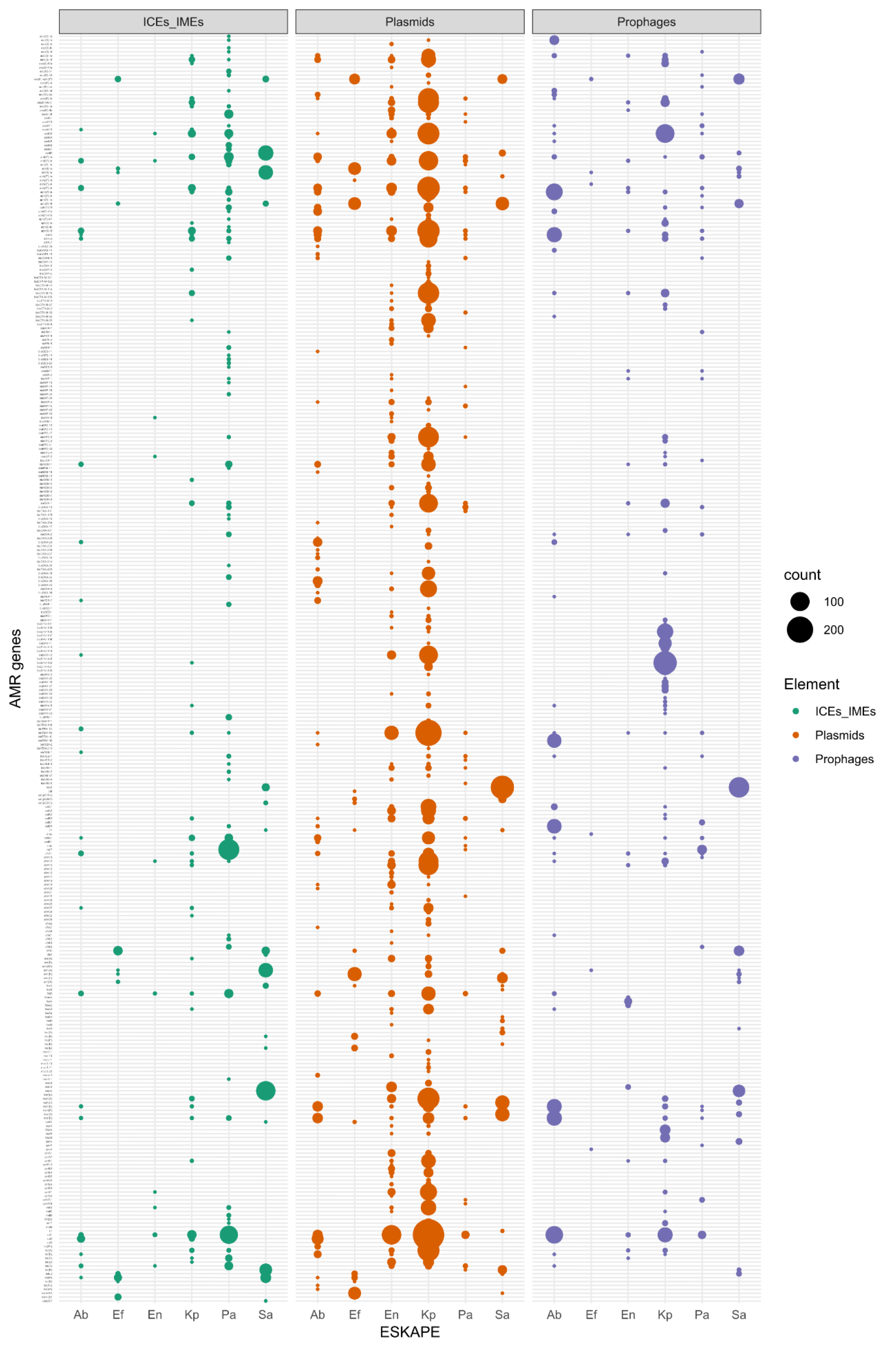
Distribution of antimicrobial resistance (AMR) genes across different ESKAPE pathogens and mobile elements. The size of the circles is proportional to absolute counts. Ab, *A. baumannii*; Ef, *E. faecium*; En, *Enterobacter* sp.; Kp, *K. pneumoniae*; Pa, *P. aeruginosa*; Sa, *S. aureus*.

**Figure S10.**
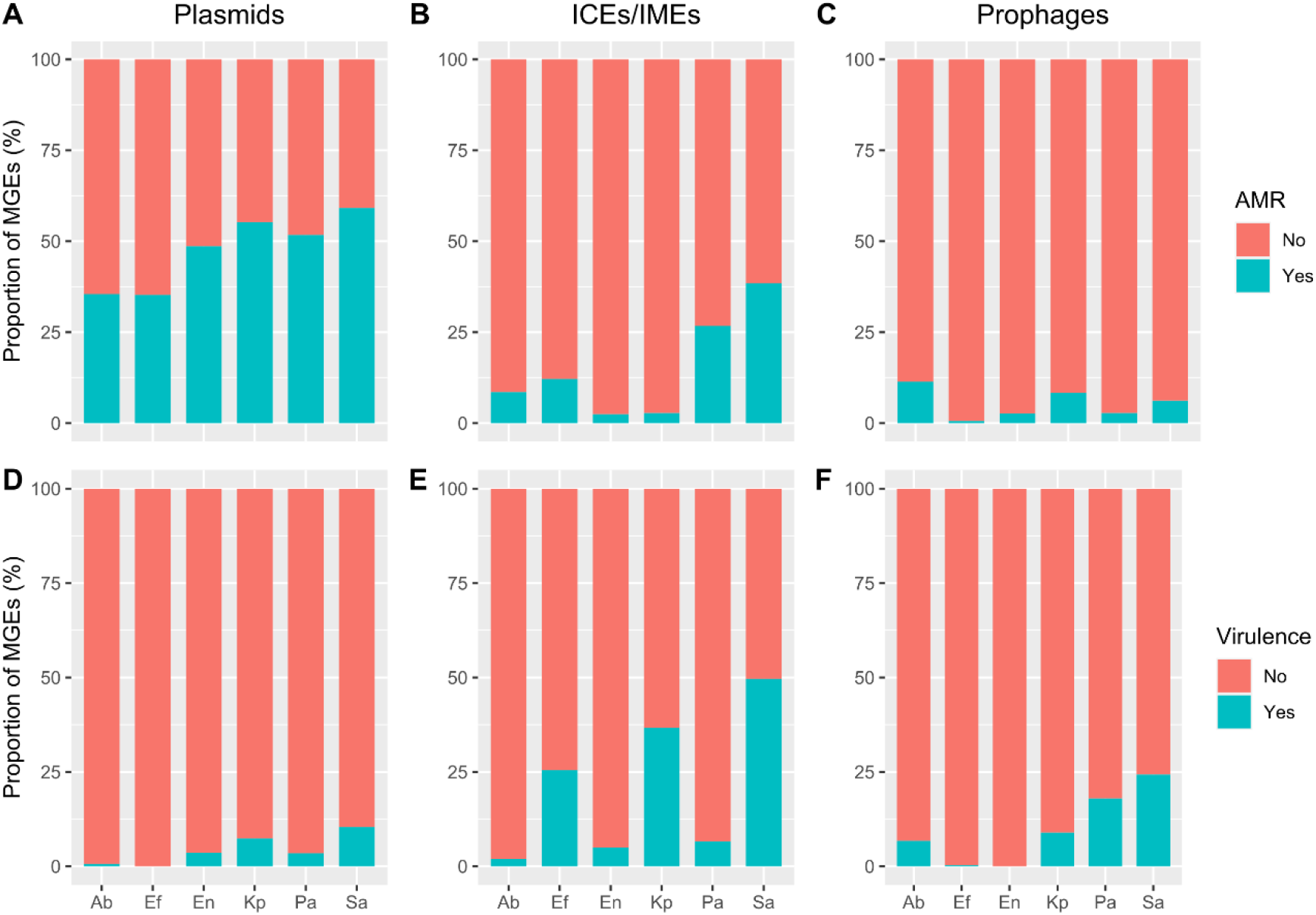
Proportion of mobile genetic elements (MGEs) carrying antimicrobial resistance (AMR) genes, identified in **A)** plasmids, **B)** ICEs/IMEs, and **C)** prophages, and virulence genes identified in **D)** plasmids, **E)** ICEs/IMEs, and **E)** prophages. A separate proportion is shown for the different ESKAPE pathogens. Each MGE was considered as positive for AMR or virulence genes if at least one of these genes were identified. Ab, *A. baumannii*; Ef, *E. faecium*; En, *Enterobacter* sp.; Kp, *K. pneumoniae*; Pa, *P. aeruginosa*; Sa, *S. aureus*.

**Figure S11.**
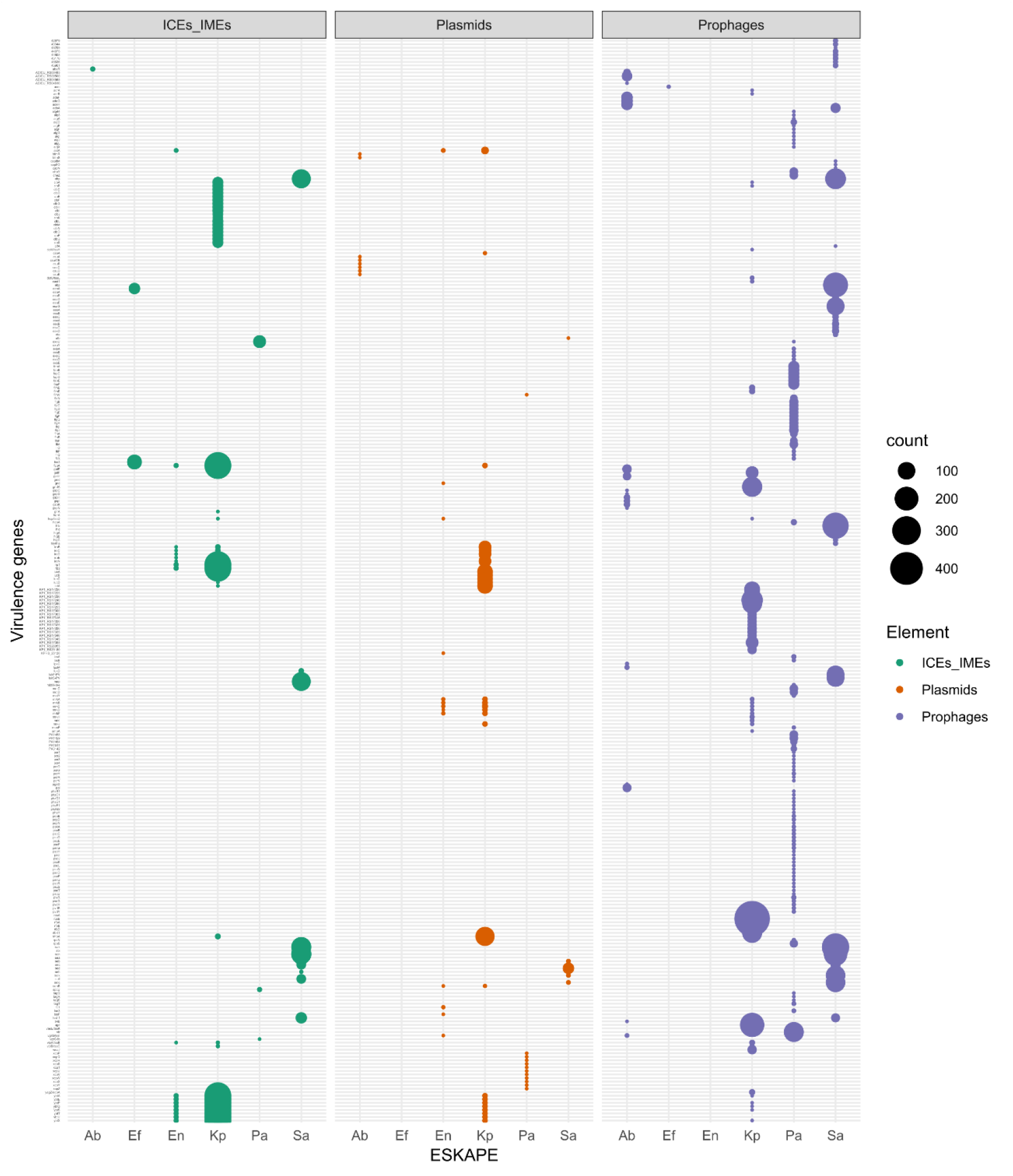
Distribution of virulence genes across different ESKAPE pathogens and mobile elements. The size of the circles is proportional to absolute counts. Ab, *A. baumannii*; Ef, *E. faecium*; En, *Enterobacter* sp.; Kp, *K. pneumoniae*; Pa, *P. aeruginosa*; Sa, *S. aureus*.

**Figure S12.**
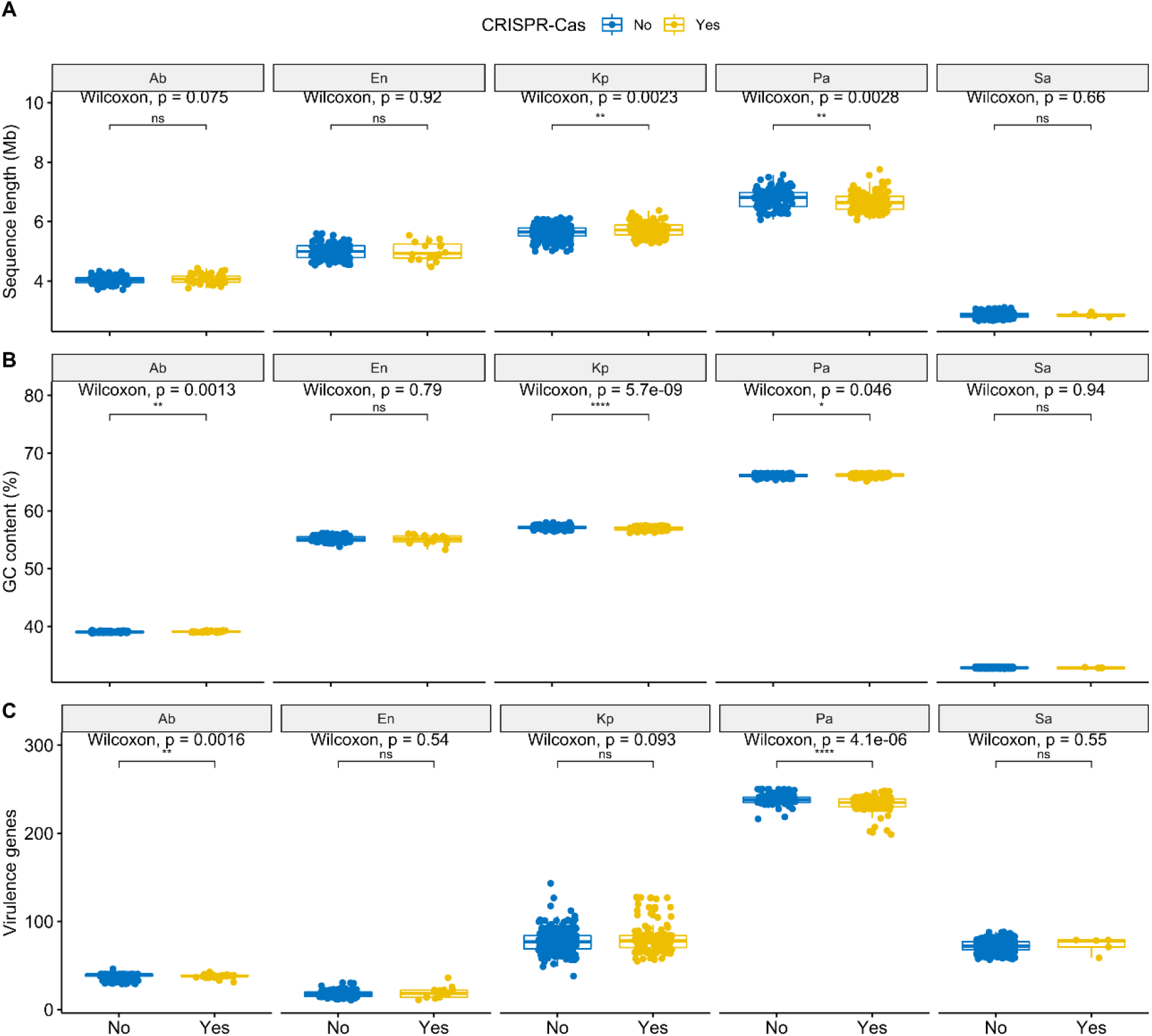
Boxplots comparing the **A)** sequence length, **B)** GC content, and **C)** the distribution of virulence genes in pairs of conspecific ESKAPE pathogens, with and without CRISPR-Cas systems. Values above 0.05 were considered as non-significant (ns). We used the following convention for symbols indicating statistical significance: * for p <= 0.05, ** for p <= 0.01, *** for p <= 0.001, and **** for p <= 0.0001. Ab, *A. baumannii*; En, *Enterobacter* sp.; Kp, *K. pneumoniae*; Pa, *P. aeruginosa*; Sa, *S. aureus*.

**Figure S13.**
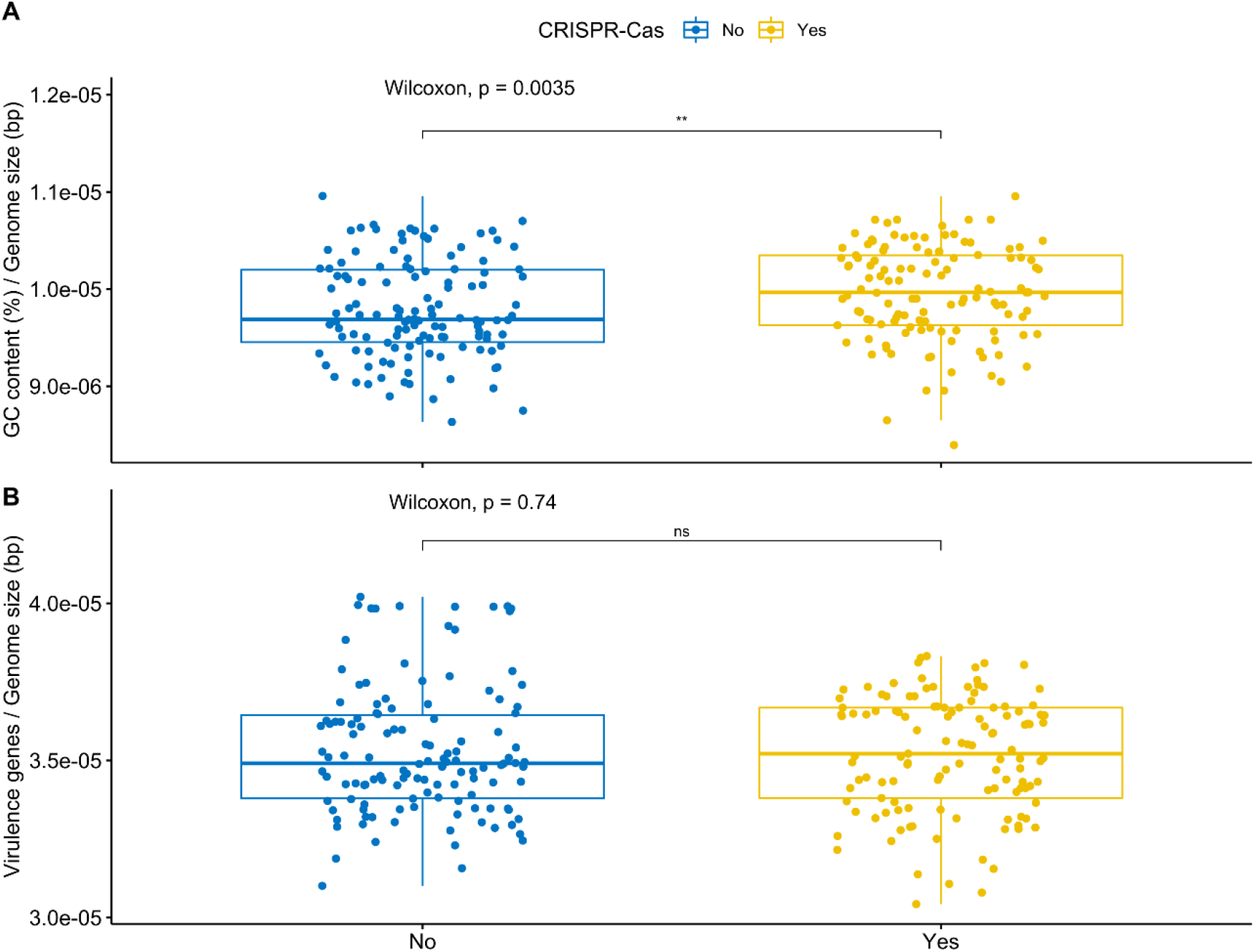
Boxplots comparing CRISPR-Cas positive and negative *P. aeruginosa* genomes, with **A)** GC content normalized to sequence length of each genome and **B)** virulence genes normalized to sequence length of each genome. Values above 0.05 were considered as non-significant (ns). We used the following convention for symbols indicating statistical significance: * for p <= 0.05, ** for p <= 0.01, *** for p <= 0.001, and **** for p <= 0.0001.

**Figure S14.**
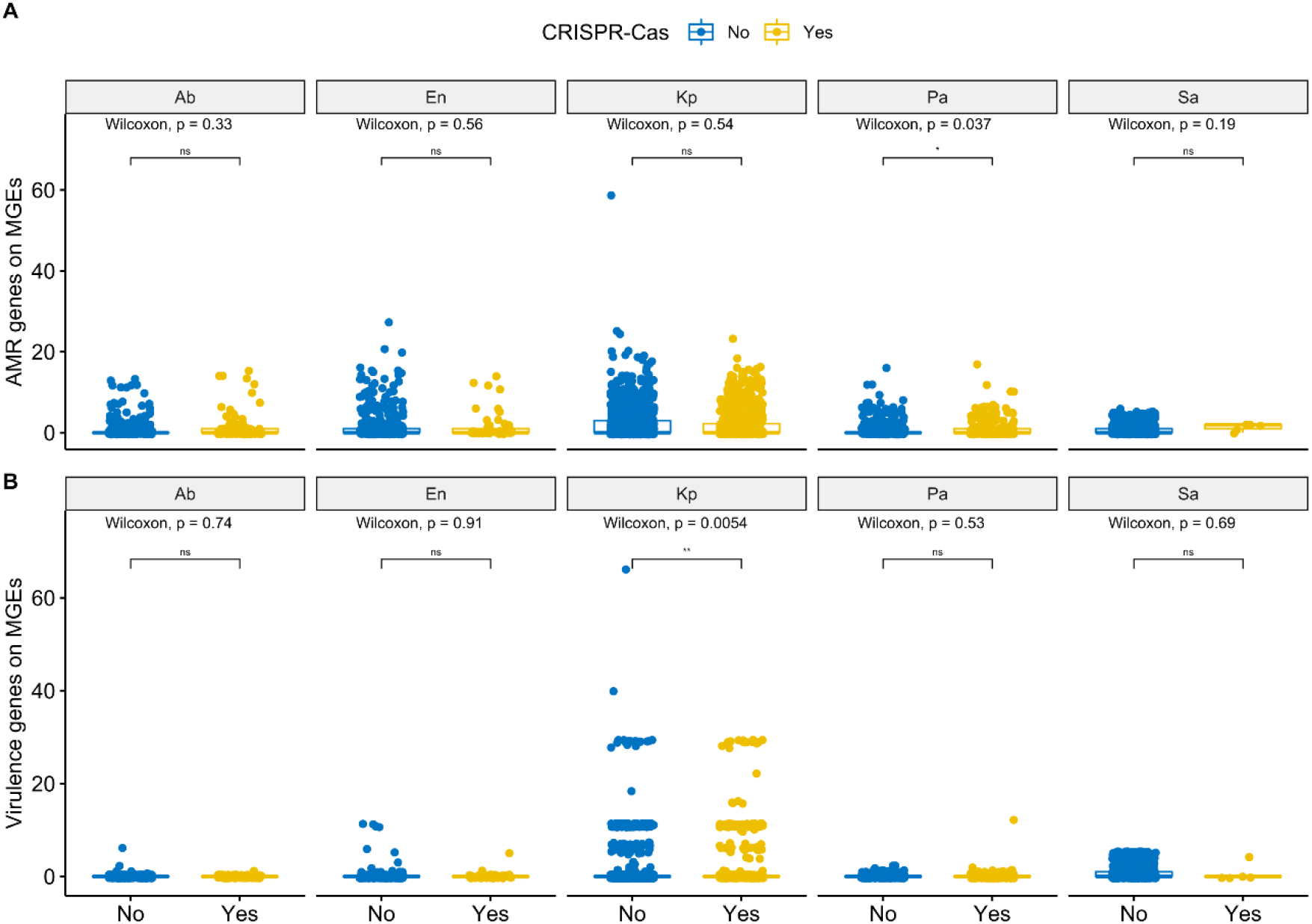
Boxplots comparing pairs of conspecific ESKAPE pathogens, with and without CRISPR-Cas systems, regarding plasmid- and ICE/IME-encoded **A**) antimicrobial resistance (AMR) and **B**) virulence genes. Values above 0.05 were considered as non-significant (ns). We used the following convention for symbols indicating statistical significance: * for p <= 0.05, ** for p <= 0.01, *** for p <= 0.001, and **** for p <= 0.0001. Ab, *A. baumannii*; En, *Enterobacter* sp.; Kp, *K. pneumoniae*; Pa, *P. aeruginosa*; Sa, *S. aureus*.

**Figure S15.**
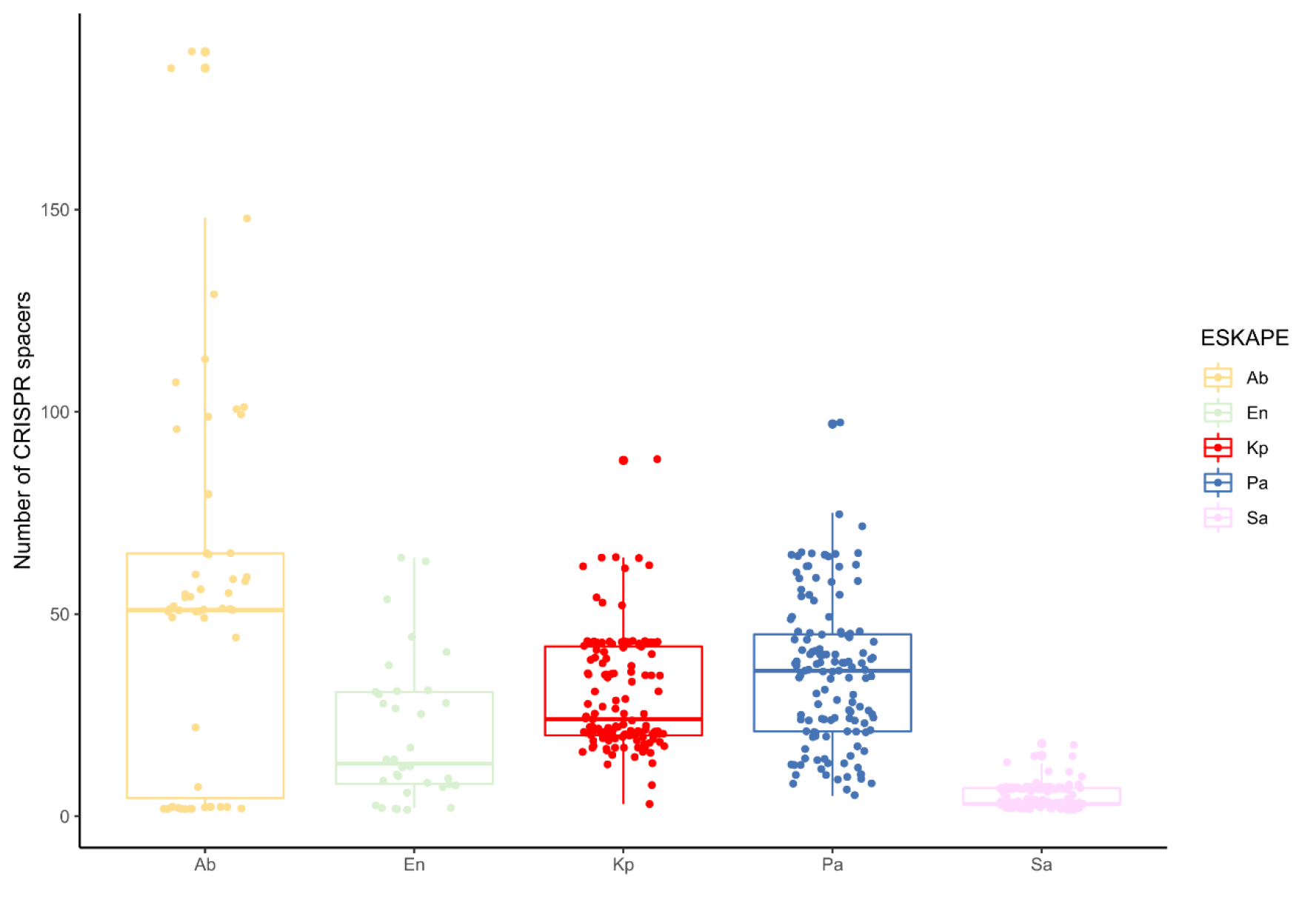
Absolute number of CRISPR spacers per masked genome identified across the ESKAPE pathogens.

## Notes

https://gitlab.gwdg.de/botelho/eskape_paper

### Competing Interest Statement

The authors have declared no competing interest.

### Summary of Updates

The identification of prophages was revised, including the methods, results, and discussion sections; figures were also revised, as well as the supplemental files.

